# Mitochondrial DNA Copy Number (mtDNA-CN) Can Influence Mortality and Cardiovascular Disease via Methylation of Nuclear DNA CpGs

**DOI:** 10.1101/673293

**Authors:** Christina A. Castellani, Ryan J. Longchamps, Jason A. Sumpter, Charles E. Newcomb, John A. Lane, Megan L. Grove, Jan Bressler, Jennifer A. Brody, James S. Floyd, Traci M. Bartz, Kent D. Taylor, Penglong Wang, Adrienne Tin, Josef Coresh, James S. Pankow, Myriam Fornage, Eliseo Guallar, Brian O’Rourke, Nathan Pankratz, Chunyu Liu, Daniel Levy, Nona Sotoodehnia, Eric Boerwinkle, Dan E. Arking

**Author notes:** corresponding author, Johns Hopkins University School of Medicine, 733 N. Broadway Miller Research Building, Room 459, Baltimore, MD, 21205.

## Abstract

**Background:** Mitochondrial DNA copy number (mtDNA-CN) has been associated with a variety of aging-related diseases, including all-cause mortality. However, the mechanism by which mtDNA-CN influences disease is not currently understood. One such mechanism may be through regulation of nuclear gene expression via the modification of nuclear DNA (nDNA) methylation.

**Methods:** To investigate this hypothesis, we assessed the relationship between mtDNA-CN and nDNA methylation in 2,507 African American (AA) and European American (EA) participants from the Atherosclerosis Risk in Communities (ARIC) study. To validate our findings we assayed an additional 2,528 participants from the Cardiovascular Health Study (CHS) (N=533) and Framingham Heart Study (FHS) (N=1,995). We further assessed the effect of experimental modification of mtDNA-CN through knockout of *TFAM*, a regulator of mtDNA replication, via CRISPR-Cas9.

**Results:** Thirty-four independent CpGs were associated with mtDNA-CN at genome-wide significance (*P*<5×10^-8^). Meta-analysis across all cohorts identified six mtDNA-CN associated CpGs at genome-wide significance (*P*<5×10^-8^). Additionally, over half of these CpGs were associated with phenotypes known to be associated with mtDNA-CN, including coronary heart disease, cardiovascular disease, and mortality. Experimental modification of mtDNA-CN demonstrated that modulation of mtDNA-CN directly drives changes in nDNA methylation and gene expression of specific CpGs and nearby transcripts. Strikingly, the ‘neuroactive ligand receptor interaction’ KEGG pathway was found to be highly overrepresented in the ARIC cohort (*P*= 5.24×10^-12^), as well as the *TFAM* knockout methylation (*P=*4.41×10^-4^) and expression (*P=*4.30×10^-4^) studies.

**Conclusions:** These results demonstrate that changes in mtDNA-CN influence nDNA methylation at specific loci and result in differential expression of specific genes that may impact human health and disease via altered cell signaling.

## BACKGROUND

Mitochondria are cytoplasmic organelles primarily responsible for cellular metabolism, and have pivotal roles in many cellular processes, including aging, apoptosis and oxidative phosphorylation[1]. Dysfunction of the mitochondria has been associated with complex disease presentation including susceptibility to disease and severity of disease[2]. Mitochondrial DNA copy number (mtDNA-CN), a measure of mitochondrial DNA (mtDNA) levels per cell, while not a direct measure of mitochondrial function, is associated with mitochondrial enzyme activity and adenosine triphosphate production. mtDNA-CN is regulated in a tissue-specific manner and in contrast to the nuclear genome, is present in multiple copies per cell, with the number being highly dependent on cell type[3, 4]. mtDNA-CN estimates can be derived from DNA isolated from blood and is therefore a relatively easily attainable biomarker of mitochondrial function. Cells with reduced mtDNA-CN show reduced expression of vital complex proteins, altered cellular morphology, and lower respiratory enzyme activity[5]. Variation in mtDNA-CN has been associated with numerous diseases and traits, including cardiovascular disease[6–8], chronic kidney disease[9], diabetes[10, 11], and liver disease[12, 13]. Lower mtDNA-CN has also been found to be associated with frailty and all-cause mortality[10].

Communication between the mitochondria and the nucleus is bi-directional and it has long been known that cross-talk between nuclear DNA (nDNA) and mtDNA is required for proper cellular functioning and homeostasis[14, 15]. Specifically, bi-directional cross-talk is essential for the maintenance and integrity of cells[16, 17], and interactions between mtDNA and nDNA contribute to a number of pathologies[18, 19]. However, the precise relationship between mtDNA and the nuclear epigenome has not been well defined despite a number of reports which have identified a relationship between mitochondria and the nuclear epigenome. For example, mtDNA polymorphisms have been previously demonstrated to be associated with nDNA methylation patterns[20] and hyper- and hypo-methylation of nuclear sites has been observed in mitochondria-depleted cancer cell lines[21]. Additionally, differential DNA methylation in brain tissue and corresponding differential gene expression were observed between strains of mice having identical nDNA, but different mtDNA[18] and reduced mtDNA-CN has been associated with inducing cancer progression via hypermethylation of nuclear DNA promoters[22]. Further, mtDNA-CN has been previously associated with changes in nuclear gene expression[23].

Thus, gene expression changes identified as a result of mitochondrial variation may be mediated, at least in part, by nDNA methylation. Further, given that it has been well-established that mtDNA-CN influences a number of human diseases we propose that one mechanism by which mtDNA-CN influences disease may be through regulation of nuclear gene expression via the modification of nDNA methylation.

To this end, we report the results of cross-sectional analysis of this association between mtDNA-CN and nDNA methylation in 5,035 individuals from the Atherosclerosis Risk in Communities (ARIC), Cardiovascular Health Study (CHS) and Framingham Heart Study (FHS) cohorts. Further, to determine the causal direction of the association between mtDNA-CN and nDNA methylation, we present results from experimental modification of mtDNA-CN followed by assessment of nDNA methylation and gene expression profiles in mtDNA-CN depleted cell lines.

## RESULTS

### mtDNA-CN is associated with nuclear DNA methylation at independent genome-wide loci in cross-sectional analysis

We performed an epigenome-wide association study (EWAS) in DNA derived from blood for 2,507 individuals from the ARIC study, comprised of 1,567 African American (AA) and 940 European American (EA) subjects (Supplemental Table S1, Supplemental Table S2). 34 independent CpGs were significantly associated with mtDNA-CN (*P*<5×10^-8^) in a meta-analysis combining the race groups (Figure 1, Supplemental Figure S1, Supplemental Table S3A) (discovery meta-analysis). This conservative *P*-value cutoff was confirmed by permutation testing. In stratified analysis of ARIC AA and EA participants, we identified 23 and 15 independent CpGs at epigenome-wide significance, respectively (Supplemental Figure S2, Supplemental Table S3B,C). Two CpGs were shared by both race groups (cg26094004 and cg21051031). ARIC AA and EA effect sizes for significant results were strongly correlated (R^2^=0.49) (Supplemental Figure S3). Further, 16/23 (70%) of AA cohort-identified CpGs showed the same direction of effect in EA participants *(P=*0.06) and 12/14 (86%) of EA cohort-identified CpGs displayed the same direction of effect in AA participants *(P=*0.008). Given these observations, we have focused on the ARIC results from combining both races (N=2,507) in further analyses. Additionally, an association was observed between increased mtDNA-CN and global hypermethylation (*P*<2.2×10^-16^, ß=0.1487) in ARIC AA, however no such association was seen in ARIC EA (*P*<0.77, ß=0.013) (Supplemental Figure S4).

**Figure 1.**
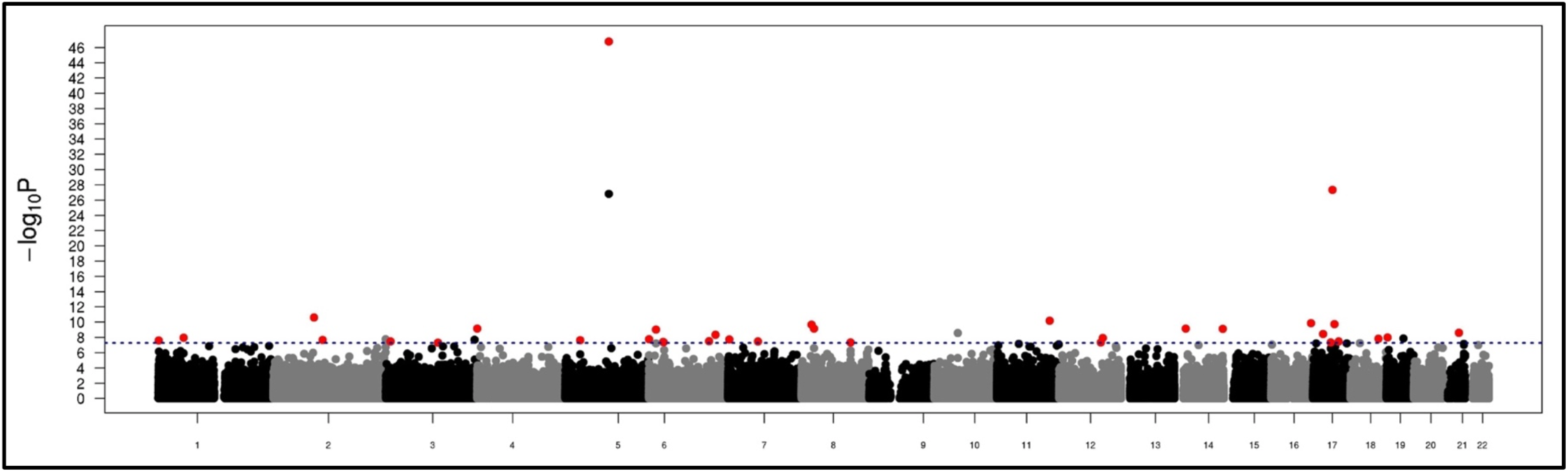
ARIC Discovery Meta-Analysis (AA and EA) Results. 34 Independent genome-wide significant CpGs were identified in ARIC meta-analysis to be associated with mtDNA-CN (red dots). Blue dotted line represents genome-wide significance cutoff (*P*=5×10^-8^). CpGs had to be independent and nominally significant in both cohorts (*P*<0.05), as well as meet the meta-analysis significance cutoff (*P*=5×10^-8^) to be considered significant.

### Pathway and biological process analysis displays associations with cell signaling functions and the ‘Neuroactive ligand-receptor interaction’ pathway

To assess the potential mechanism underlying the identified associations we performed GO and KEGG pathway analysis. mtDNA-CN associated CpGs were annotated with their nearest gene. KEGG analysis identified the *neuroactive ligand-receptor interaction pathway (path:hsa04080)* to be the top overrepresented pathway (*P=* 5.24×10^-12^, Permuted *P*=3.84×10^-5^) (Table 1a). Further, GO analyses identified a number of biological processes related to cell signaling and ligand interactions which included *Cell-cell signaling* (*P*=1.42×10^-3^)*, Trans-synaptic signaling* (*P*=1.88×10^-3^) *and Synaptic signaling* (*P*=1.88×10^-3^), among others (Table 1b). These results met permutation tested *P*-value cutoffs.

**Table 1.** Results of pathway and functional analysis in ARIC discovery meta-analysis and *TFAM* knockout methylation and expression datasets with combined *P-Value* and ARIC *P-Value* <0.05. *TFAM* integrated (INT) *P*-value represents combined methylation and expression results. Combined *P*-value represents combined ARIC and *TFAM* methylation and expression results. **A.** KEGG pathways sorted by combined p-value. **B.** GO pathways sorted by combined p-value.

| Pathway | Name | ARIC<br>*P*-value | *TFAM*<br>METH<br>*P*-value | *TFAM*<br>RNA<br>*P*-value | *TFAM*<br>INT<br>*P*-value | Combined Pathway *P*-value<br>(ARIC, *TFAM*-METH,<br>*TFAM*-RNA) |
| --- | --- | --- | --- | --- | --- | --- |
| path:hsa04080 | Neuroactive ligand-receptor interaction | 5.24E-12 | 4.41E-04 | 4.30E-04 | 8.77E-06 | 8.96E-16 |
| path:hsa05033 | Nicotine addiction | 8.99E-04 | 6.30E-05 | 9.32E-04 | 7.72E-06 | 1.61E-08 |
| path:hsa04024 | cAMP signaling pathway | 1.29E-05 | 2.32E-02 | 2.23E-01 | 3.25E-01 | 1.03E-05 |
| path:hsa04614 | Renin-angiotensin system | 1.04E-05 | 5.11E-02 | 1.00E+00 | 1.00E+00 | 6.37E-05 |
| path:hsa04723 | Retrograde endocannabinoid signaling | 1.89E-04 | 1.84E-02 | 1.56E-01 | 3.66E-03 | 6.48E-05 |
| path:hsa05032 | Morphine addiction | 1.26E-02 | 4.38E-02 | 5.15E-02 | 1.40E-03 | 1.88E-03 |
| path:hsa05031 | Amphetamine addiction | 3.54E-02 | 9.64E-02 | 4.23E-01 | 1.19E-01 | 4.18E-02 |
| path:hsa04724 | Glutamatergic synapse | 1.80E-02 | 9.49E-02 | 1.00E+00 | 1.00E+00 | 4.73E-02 |

**B.**
| Function | Name | ARIC<br>*P*-value | *TFAM*<br>METH<br>*P*-value | *TFAM*<br>RNA<br>*P*-value | *TFAM*<br>INT<br>*P*-value | Combined Function *P*-value<br>(ARIC, *TFAM*-METH,<br>*TFAM*-RNA) |
| --- | --- | --- | --- | --- | --- | --- |
| GO:0007267 | Cell-cell signaling | 1.42E-03 | 1.71E-05 | 1.19E-02 | 1.42E-02 | 7.63E-08 |
| GO:0099537 | Trans-synaptic signaling | 1.88E-03 | 1.06E-05 | 6.27E-02 | 1.92E-02 | 2.89E-07 |
| GO:0099536 | Synaptic signaling | 1.88E-03 | 1.08E-05 | 6.34E-02 | 1.92E-02 | 2.97E-07 |
| GO:0007268 | Chemical synaptic transmission | 1.88E-03 | 2.22E-05 | 6.05E-02 | 1.87E-02 | 5.47E-07 |
| GO:0098916 | Anterograde trans-synaptic signaling | 1.88E-03 | 2.22E-05 | 6.05E-02 | 1.87E-02 | 5.47E-07 |
| GO:0099095 | Ligand-gated anion channel activity | 4.30E-02 | 8.02E-04 | 1.23E-04 | 3.74E-07 | 8.74E-07 |
| GO:0045202 | Synapse | 1.98E-04 | 9.81E-05 | 2.74E-01 | 5.92E-03 | 1.07E-06 |
| GO:0045211 | Postsynaptic membrane | 8.33E-03 | 1.03E-04 | 1.45E-01 | 6.19E-04 | 1.78E-05 |

### Validation of CpG associations in independent cohorts

We performed a validation study to replicate findings from the ARIC discovery population in blood samples from 239 AA and 294 EA participants from CHS as well as 1,995 EA participants from FHS, for a total of 2,528 individuals (Supplemental Table S1). 7/34 CpGs identified in the discovery cohort were nominally significant (*P*<0.05 and same direction of effect as the ARIC cohort results) (Supplemental Table S3) and the effect sizes from the ARIC results and the validation meta-analysis were largely correlated (R^2^=0.36) (Figure 2). Overall, the results were consistent across individual cohorts (Supplemental Figure S5, Supplemental Table S3) and analysis of the results from the 34 CpGs across all three cohorts (ARIC, CHS and FHS, N=5,035), identify six CpGs as validated mtDNA-CN associated CpGs (*P*<5×10^-8^) (Supplemental Table S3, Supplemental Figure S6).

**Figure 2.**
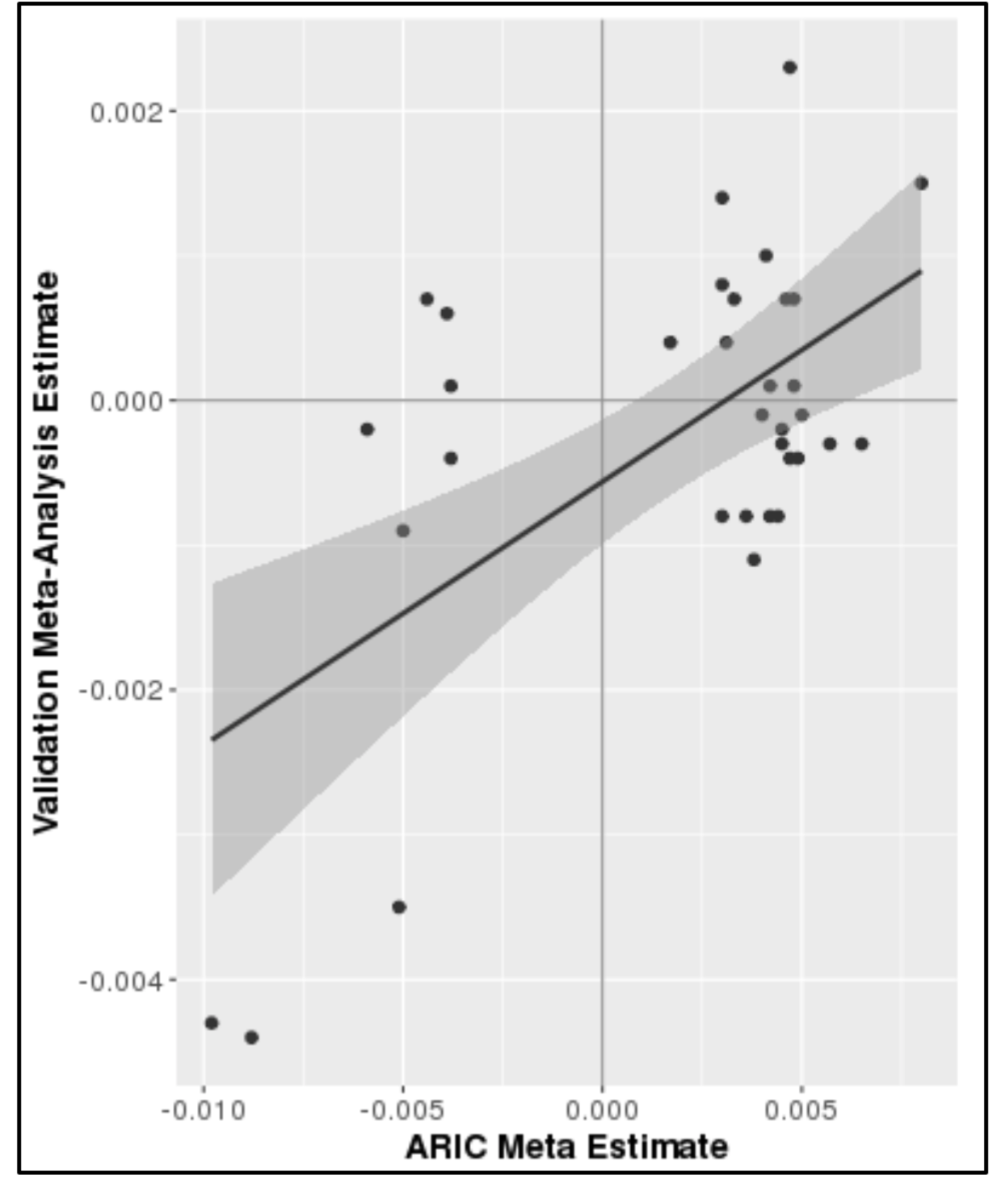
Validation of Meta-analysis identified CpGs in CHS and FHS combined cohorts (N=2,528, R^2^=0.36, Kendall tau=0.19).

### Establishing causality via TFAM knockout: Pathway and biological process analysis of TFAM KO methylation and expression results independently identify pathways observed in cross-sectional analysis

To assess if modification of mtDNA-CN drives changes to nuclear DNA methylation we used CRISPR-Cas9 to knock out the *TFAM* gene, which encodes a regulator of mtDNA replication (Figure 3). Knockout of *TFAM* has been shown to reduce mtDNA-CN[24, 25]. Heterozygous knockout of the *TFAM* gene (confirmed by qPCR of *TFAM* DNA) in HEK293T cells resulted in a 5-fold reduction in the steady-state expression levels of *TFAM*, a marked reduction in protein production (>81%), and an 18-fold reduction in mtDNA-CN across three independent knockout events (Figure 4, Supplemental Figure S7). We assayed methylation and expression of genes in the three knockout lines using the Illumina Infinium Methylation EPIC Beadchip and RNA-sequencing (Supplemental Table S4, Supplemental Table S5).

**Figure 3.**
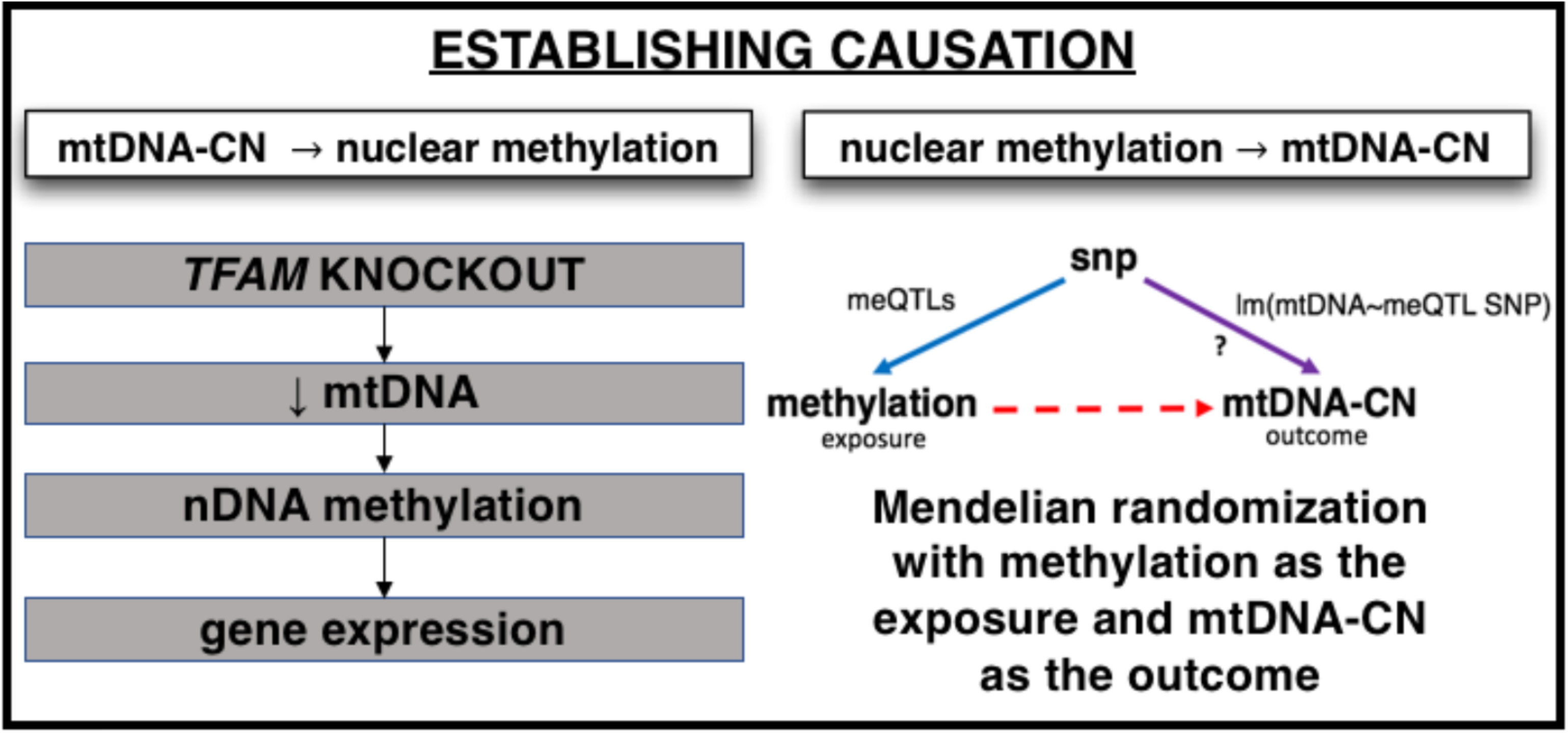
Methods used to establish causation.

**Figure 4.**
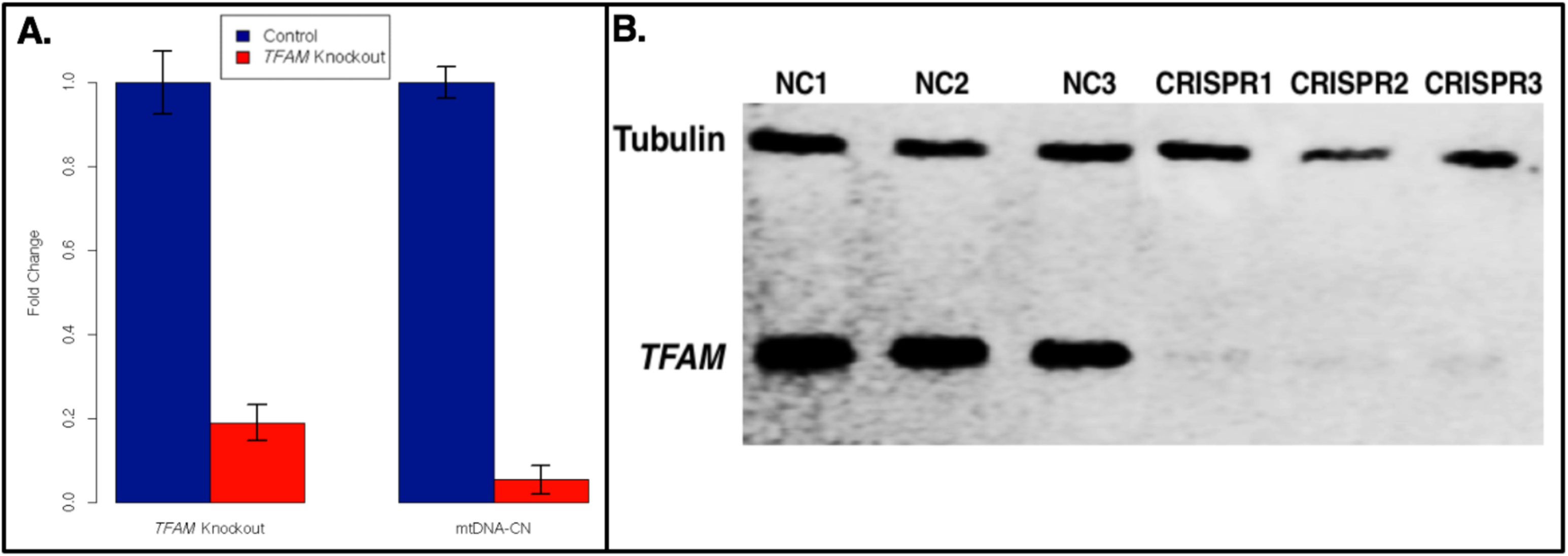
CRISPR-Cas9 induced heterozygous knockout of *TFAM* reduced RNA expression, mtDNA-CN and protein levels. **A.** RNA expression was reduced by over 80% relative to negative control (NC) expression (left) (passage 45). mtDNA-CN levels showed an ∼18-fold reduction in *TFAM* knockout cell lines, (passage 32) (right). **B.** Western blot of CRISPR *TFAM* heterozygous knockout showed a significant reduction (>81%) in *TFAM* protein (passage 35). NC=Negative Control lines. CRISPR=CRISPR *TFAM* knockout lines. Control is Tubulin.

We hypothesized that if similar mechanisms are at play in blood and kidney then consistent pathways would be identified between the cross-sectional cohort analysis and our *TFAM* KO analysis. We therefore identified over-represented terms resulting from GO and KEGG analysis of differentially methylated CpGs and differentially expressed genes as well as gene-level integrated methylation and expression results (cutoffs used: *TFAM* Methylation - top 300 differentially methylated CpGs, *TFAM* Expression – differentially expressed genes (169 genes), *TFAM* Integrated Methylation/Expression – top 188 genes). The TFAM KO pathway analysis results are consistent with the findings from our ARIC cross-sectional analysis. Specifically, KEGG analysis identified the *neuroactive ligand-receptor interaction pathway (path:hsa04080)* to be the second most overrepresented pathway in the *TFAM* knockout methylation analysis (*P=*4.41×10^-4^) and the top overrepresented pathway in the *TFAM* knockout RNA sequencing analysis (*P=*4.30×10^-4^) (Table 1a). Accordingly, integration of results from *TFAM* knockout methylation and expression also resulted in strong association with this pathway (*P=*8.77×10^-6^). Further, combining of *P*-values (Fisher’s method) across the ARIC meta-analysis, *TFAM* knockout methylation and *TFAM* knockout expression analyses yielded a combined *P*-value of 8.96×10^-16^ for this pathway which was also the top pathway identified in integrated analysis (Table 1a).

The specific genes identified by each analysis to be part of the *neuroactive ligand receptor interaction pathway* were unique to each study (Supplemental Table S6), with only one gene (*GABRG3*) in common between ARIC analyses and *TFAM* knockout methylation analysis and only one gene (*GABRB1*) in common between *TFAM* knockout methylation and expression analyses (Supplemental Table S6).

GO analyses of *TFAM* knockout cell lines also confirmed the finding from cross-sectional analysis that biological processes related to cell signaling and ligand interactions including *Cell-cell signaling* (combined *P*=7.63×10^-8^)*, Trans-synaptic signaling* (combined *P*=2.89×10^-7^) *and Synaptic signaling* (combined *P*=2.97×10^-7^) were over-represented, among others (Table 1b). These results suggest that mtDNA-CN drives changes to nDNA methylation at sites nearby genes relating to cell signaling processes which in turn may cause gene expression changes to these genes and contribute to disease.

#### mtDNA-CN is causative of changes in nuclear DNA methylation and nuclear gene expression

Although we do not *a priori* expect site-specific methylation results to be consistent with those we have identified in blood due to the fact that HEK293T lines represent a different tissue and are subject to the inherent variability of cell culture systems, we note that three validated mtDNA-CN associated CpGs showed nominally significant differential expression (*P*<0.05) and two were significant after Bonferroni correction (*P*<0.01, Table S7, Supplemental Figure S8). We also observe no difference in global methylation patterns between negative control and *TFAM* knockout cell lines (Supplemental Figure S9). All nominally differentially expressed genes (*P*<0.05) within 1Mb of the *TFAM* knockout differentially methylated CpGs were identified (Supplemental Table S8). Five genes nearby the three differentially methylated CpGs were differentially expressed after Bonferroni correction for the number of genes within 1Mb of each CpG (*P*<6.41×10^-4^) (Table 2). The five differentially expressed genes were: *IFI35* (*P*=3.76×10^-5^) and *RAMP2* (*P*=5.51×10^-4^) near cg26094004; *RPIA* near cg26563141 (*P*=5.04×10^-6^); and *HLA-DRB5* (*P*=6.50×10^-7^) and *MSH5* (*P*=2.50×10^-4^) near cg08899667.

**Table 2.**
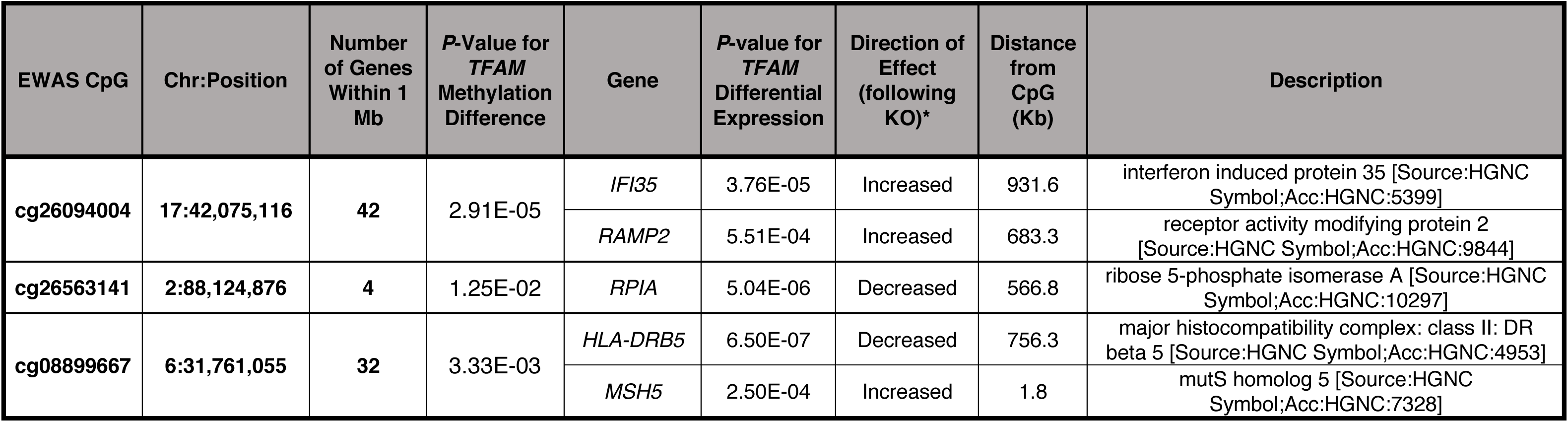
Differentially expressed genes (*P*=6.41×10^-4^) within 1Mb of differentially methylated CpGs.

### Establishing causality via Mendelian Randomization (MR): Nuclear DNA methylation does not appear to be causative of changes in mtDNA-CN at identified CpGs

Mendelian randomization (MR), a form of instrument variable analysis, is a well-established method for the assessment of causal relationships. Specifically, MR seeks to establish causality by exploiting the fact that SNPs are assigned at conception and randomly distributed in the population making them an excellent instrument variable to determine the relationship between modifiable exposures and relevant outcomes. We used MR to further test the direction of causality between mtDNA-CN (outcome) and nuclear methylation (exposure) by exploring the relationship between methylation quantitative trait loci (meQTLs) (instrument variable) and mtDNA-CN (Figure 3, Supplemental Table S9). Specifically, if nDNA methylation at our sites of interest is causative of changes in mtDNA-CN, then meQTL single nucleotide polymorphisms (SNPs) for these CpGs of interest would be expected to also be associated with mtDNA-CN. Alternatively, if mtDNA-CN is not associated with meQTL SNPs, then it would follow that changes to nDNA methylation likely do not drive changes to mtDNA-CN at these CpGs, or suggest that other factors are modulating these interactions.

We identified four independent *cis* meQTLs in the ARIC EA cohort (Permuted *P*=7.84×10^-4^) and six independent *cis* meQTLs in the ARIC AA cohort (Permuted *P*=9.12×10^-4^) across five mtDNA-CN associated CpGs for use as an instrument variable for MR (Supplemental Table S9A). We further identified two independent discovery meta-analysis derived meQTLs by combining results from ARIC EA and AA cohorts (Permuted *P*=3.97×10^-5^, fixed effects (FE) model) (Supplemental Table S9B).

We then assessed the relationship between meQTL SNPs and mtDNA-CN. The results of the MR were null for each independent meQTL (Bonferroni *P*=0.005) (Supplemental Table S9). While our power for a single meQTL varied depending on the specific meQTL assessed, with power to detect an individual association ranging from 0.18 to 0.99 across the 12 meQTLs, overall power was >99% to detect at least one associated meQTL. These results support the experimentally established direction of causality by suggesting that modification of nDNA methylation at CpG sites of interest does not directly drive alterations in mtDNA-CN.

### Association of CpG methylation with mtDNA-CN associated phenotypes

Since decreased mtDNA-CN has been associated with a number of aging-related diseases, and given our hypothesis that mtDNA-CN leads to nDNA methylation changes which influence disease outcomes, associated CpGs should also be associated with mtDNA-CN related phenotypes. To test these associations, we performed linear regression and survival analysis for prevalent and incident diseases, respectively, for each of the six validated CpGs as they relate to coronary heart disease (CHD), cardiovascular disease (CVD), and mortality in the ARIC, FHS and CHS cohorts (Table 3, Supplemental Table S10). Results from each cohort were meta-analyzed to derive an overall association for each validated CpG with each outcome of interest.

**Table 3.** Summary of phenotype associations from all-cohort meta-analysis for Validated CpGs. Bold entries highlight nominally significant associations (*P*<0.05). *Significant after stringent multiple test correction.

| All Cohorts | Meta Direction | Expected Direction of Phenotype Association | Prevalent CHD |  |  | Prevalent CVD |  |  | Incident CHD |  |  | Incident CVD |  |  | Mortality |  |  |
| --- | --- | --- | --- | --- | --- | --- | --- | --- | --- | --- | --- | --- | --- | --- | --- | --- | --- |
|  |  |  | Beta | Std. Error | P-Value | Beta | Std. Error | P-Value | Beta | Std. Error | P-Value | Beta | Std. Error | P-Value | Beta | Std. Error | P-Value |
| cg21051031 | Positive | Negative | -0.56 | 1.95 | 7.74E-01 | -0.67 | 1.88 | 7.23E-01 | 0.96 | 1.80 | 5.94E-01 | -0.28 | 1.41 | 8.43E-01 | -0.79 | 0.81 | 3.31E-01 |
| cg26094004 | Negative | Positive | 2.91 | 1.48 | **4.89E-02** | 3.13 | 1.40 | **2.55E-02** | 0.89 | 0.93 | 3.39E-01 | -0.13 | 0.77 | 8.63E-01 | -0.78 | 0.49 | 1.16E-01 |
| cg26563141 | Negative | Positive | 0.22 | 1.33 | 8.69E-01 | -0.31 | 1.24 | 8.00E-01 | 0.13 | 0.94 | 8.88E-01 | 1.10 | 0.75 | 1.40E-01 | 0.98 | 0.45 | **2.87E-02** |
| cg14575356 | Positive | Negative | -1.80 | 2.04 | 3.79E-01 | -2.19 | 1.88 | 2.45E-01 | 2.71 | 1.34 | **4.36E-02** | 1.75 | 1.08 | 1.04E-01 | 0.12 | 0.68 | 8.63E-01 |
| cg23513930 | Positive | Negative | 1.99 | 3.37 | 5.54E-01 | 0.88 | 3.07 | 7.75E-01 | 2.24 | 1.84 | 2.24E-01 | 0.50 | 1.49 | 7.39E-01 | 1.07 | 0.94 | 2.53E-01 |
| cg08899667 | Negative | Positive | 3.65 | 1.76 | **3.81E-02** | 3.35 | 1.61 | **3.81E-02** | 1.57 | 1.10 | 1.54E-01 | 0.06 | 0.93 | 9.44E-01 | 2.08 | 0.59 | **3.93E-04\*** |

We identify nominally significant phenotype associations with at least one of the mtDNA-CN associated traits of interest for four of the six validated mtDNA-CN associated CpGs (*P*<0.05). Specifically, results in the expected direction of effect for prevalent CHD and prevalent CVD were identified for two mtDNA-CN associated CpGs (cg26094004 and cg08899667). Similarly, results in the expected direction of effect were identified for the association between all-cause mortality and cg26563141 and cg08899667. Thus, we found cg08899667 to be nominally associated with three of the five mtDNA-CN associated phenotypes, including all-cause mortality (Table 3).

## DISCUSSION

We report evidence that changes in mtDNA-CN influence nDNA methylation at specific, validated loci, including those acting in the ‘neuroactive ligand receptor interaction’ pathway which may impact human health and disease via altered cell signaling. A number of these associations were validated across two independent cohorts and identified both cross-sectionally and experimentally. It is important to note that the methods used to estimate mtDNA-CN differed between the three cohorts with a qPCR based approach used for CHS, a whole-genome sequencing approach for FHS and microarray analysis for ARIC. This may reflect the robustness of results across mtDNA-CN estimation methods and also explain why some but not all CpGs replicated in our validation analysis[26]. We also note that as expected, our experimental approach using kidney cell lines replicated some but not all of the pathway-level results from blood. These findings likely reflect both the intrinsic differences between cell line data and cross-sectional data as well as the inherent complexity of mitochondrial-to-nuclear signaling which would be expected to vary across cell-types, developmental timepoints and environmental conditions.

### DNA methylation as a link between mtDNA-CN and changes in nuclear gene expression

The relationship between the nuclear and mitochondrial genomes strongly implicates communication between them as vital for proper cell functioning. Given the function of the mitochondria in meeting cellular energy demands, mitochondria may play an important role in translating environmental stimuli into epigenetic changes. Accordingly, mtDNA-CN levels are sensitive to a number of chemicals[27], highlighting the role of mtDNA as an environmental biosensor. We hypothesize that modification of nDNA methylation via mitochondrial signaling modifies gene expression which in turn may lead to disease outcomes or influence severity of disease. Supporting this, epigenetic changes in nuclear DNA correlate with reduced cancer survival and low mtDNA-CN correlates with poor survival across a number of cancer types[28, 29]. Thus, retrograde signals from the mitochondria to the nucleus may be crucial in sensing homeostasis and translating extracellular signals into altered gene expression[18].

Our results implicate the *neuroactive ligand receptor interaction* pathway and in general additional processes involved in cellular signaling. The results also show that although the same pathways are implicated across our independent datasets, the specific genes affected differ between conditions. Interestingly, the *neuroactive ligand receptor interaction* pathway has been identified as having the second highest number of atherosclerosis candidate genes of any KEGG pathway, harboring 53 atherosclerosis candidate genes (272 total genes in the pathway)[30]. This is an interesting finding given the association of mtDNA-CN with cardiovascular disease[6–8]. Perhaps unsurprisingly, this pathway also belongs to the class of KEGG pathways that are responsible for environmental information processing and signaling molecules/interactions. Genes from this pathway are highly expressed across a wide variety of tissues including whole blood, heart tissue and cardia myocytes, among others (http://software.broadinstitute.org/gsea/). We further observe synaptic signaling to be significantly associated, which is supported by observations that synaptic signals can induce changes in nDNA methylation leading to plasticity related gene expression changes[31].

### Proposed mechanisms for the methylation of nDNA as a result of changes in mtDNA

The precise identity of the signal(s) coming from the mitochondria that might be responsible for modifying nDNA methylation has not yet been identified and warrants further experimentation. It is however possible that histones may play a role in this signaling process. Supporting this, mitochondria-to-nucleus retrograde signaling has been shown to regulate histone acetylation and alter nuclear gene expression through the heterogenous ribonucleoprotein A2 (hnRNAP2)[23]. In fact, histone modifications co-vary with mitochondrial content and are linked with chromatin activation, namely H4K16, H3L4me3 and H3K36me2[32]. In addition, perturbations to oxidative phosphorylation alter methylation processes by modifications to the methionine cycle. Methionine metabolism is essential for production of *S*-adenosylmethioninine (SAM), a methyl donor for histone and DNA methyltransferases[33].

Uncovering the precise nature of this signaling from mitochondria to the nucleus would be expected to expose essential clues that will integrate epigenetic regulation, mitochondrial and genomic polymorphisms, and complex phenotypes. Further assessment of the functional mechanisms underlying the crosstalk between mtDNA-CN, methylation and disease through sophisticated cell-culture models will be required to fully appreciate the diagnostic and therapeutic utility of the interaction between mtDNA and nDNA as identified in this study.

### Influence of findings on complex disease etiology

The observation that differential methylation occurred at specific-sites throughout the nuclear genome as a result of changes to mtDNA-CN, provides an explanation for how mtDNA could alter normal homeostasis as well as susceptibility and/or severity of diseases. The association of mtDNA-CN associated CpGs with mtDNA-CN related disease states lends further support to the hypothesis that modulation of mtDNA-CN not only modifies the nuclear epigenome, and the expression of nearby genes, but does so at locations which may be relevant to disease outcomes, including cardiovascular disease and all-cause mortality. In particular, these observations may explain how mitochondrial-to-nuclear signaling could influence polygenic traits with complex etiology and in particular those for which environmental insults play a role.

Further, these findings have direct implications for the recent emergence of mitochondrial donation in humans as they suggest that mitochondrial replacement into recipient oocytes may lead to unexpected changes to the nuclear epigenome. Thus, unravelling the complex interplay of the mitochondria and nucleus is also critical to properly informing medical decision makers.

This study design had a number of strengths and limitations. A possible limitation of the cross-sectional analysis is the potential for some common factor we have not been able to account for to influence both mtDNA-CN and nDNA methylation. We also note that future analyses in larger admixed cohorts can identify and validate race-specific loci of interest. In experimental analysis, we used HEK293T cells for our knockdown studies due to the optimized protocols available as well as their propensity for transfection; however, we note that the use of a blood cell line may be more relevant to direct interpretation of the results. We also note that the *TFAM* knockout may independently influence nDNA methylation though unknown mechanisms that we did not account for in this study and/or *TFAM* may have an effect on differentially expressed genes that is independent of the epigenome. Additionally, given that transcriptional activation/repression can be regulated over large distances in the genome, our approach using the nearest gene for pathway analysis may have limited the number of associations identified. Further, prevalent disease is subject to reverse causality and therefore the results on prevalent phenotypes should be interpreted with caution. Strengths of this study include the well phenotyped and carefully collected incident disease data, the robustness of the findings across multiple cohorts and ethnic groups, as well as the careful quality control employed. Further, our results stood up to rigorous permutation testing which increases the reliability of these observations.

## CONCLUSIONS

Cross-sectionally we have shown that variation in mtDNA-CN is associated with nuclear epigenetic modifications at specific CpGs across multiple independent cohorts. Specifically, six mtDNA-CN associated CpGs were robustly identified across three independent cohorts. Second, we found meQTL SNPs to not be associated with mtDNA-CN, suggesting that nuclear methylation at these CpGs does not directly cause altered mtDNA-CN. Third, functional results show that modulation of mtDNA-CN leads to differential methylation and expression of genes relating to cell signaling processes. Further, mtDNA-CN associated CpGs display association with mtDNA-CN related phenotypes, namely cardiovascular disease and all-cause mortality. These findings demonstrate that the mechanism(s) by which mtDNA-CN influences disease is at least in part via regulation of nuclear gene expression through modification of nDNA methylation. Specifically, the data presented here support the model that modification of mtDNA-CN leads to changes to nDNA methylation which in turn influence nuclear DNA expression of nearby genes which contribute to disease pathology. These results have implications for understanding the mechanisms behind mitochondrial and nuclear communication as it relates to complex disease etiology as well as the consequences of mitochondrial replacement therapeutic strategies. Taken together, the results confirm that in elucidating the underpinnings of complex disease, knowledge of only nuclear DNA dynamics is not sufficient to fully elucidating disease etiology.

## MATERIALS AND METHODS

### Ethics

All studies were approved by their respective Institutional Review Boards (see supplemental methods).

### Discovery Study Analysis

#### The Atherosclerosis Risk in Communities Cohort (ARIC)

The ARIC study is a prospective cohort intended for the study of cardiovascular disease in subjects from four communities across the USA: Forsyth County, NC, northwest suburbs of Minneapolis, MN, Jackson, MS, and Washington County, MD[34]. Sample characteristics are available in Supplemental Table S1. Following quality control, 1,567 African Americans (AA) and 940 European Americans (EA) were used as a discovery cohort. Participants for ARIC EA were derived from two existing projects, Brain MRI (81.7%) and OMICS (18.3%). DNA was extracted from peripheral blood leukocyte samples from visit 2 or 3 using the Gentra Puregene Blood Kit (Qiagen; Valencia, CA, USA) according to the manufacturer’s instructions (www.qiagen.com) and hybridized to the Illumina Infinium Human Methylation 450K BeadChip and the Genome-Wide Human SNP Array 6.0.

#### Estimation of mtDNA-CN from Affymetrix Human SNP 6.0 Arrays

The Affymetrix Genome-Wide Human SNP 6.0 Array was used to estimate mtDNA-CN for each participant as previously described[35]. Briefly, mtDNA copy number (mtDNA-CN) was determined utilizing the Genvisis software package (http://www.genvisis.org). Initially, a list of high-quality mitochondrial SNPs were hand-curated by employing BLAST to remove SNPs without a perfect match to the annotated mitochondrial location and SNPs with off-target matches longer than 20 bp. The probe intensities of the 25 remaining mitochondrial SNPs was determined using quantile sketch normalization (apt-probeset-summarize) as implemented in the Affymetrix Power Tools software. To correct for DNA quality, DNA quantity, hybridization efficiency and other technical artifacts, surrogate variable analysis was applied to the BLAST filtered, GC corrected log R ratio (LRR) of 43,316 autosomal SNPs. These autosomal SNPs were selected based on the following quality filters: call rate >98%, HWE *P*-value >0.00001, PLINK mishap for non-random missingness *P*-value >0.0001, association with sex *P*-value 0.00001, linkage disequilibrium pruning (r^2^ <0.30), maximal autosomal spacing of 41.7 kb. The median of the normalized intensity, LRR for all homozygous calls was GC corrected and used as initial estimates of mtDNA-CN for each sample. The final measure of mtDNA-CN is represented as the standardized residuals from a race-stratified linear regression adjusting the initial estimate of mtDNA-CN for 15 surrogate variables (SVs), age, sex, sample collection site, and white blood cell count. Technical covariates such as DNA quality, DNA quantity, and hybridization efficiency were captured via surrogate variable analysis (SVA) as previously described[7, 36].

#### Illumina Infinium Human Methylation 450K Beadchip Analysis

The Infinium Human Methylation 450K BeadChip was used to determine DNA methylation profiles from blood for >450,000 CpGs across the human genome.

##### Bisulfite Conversion

Bisulfite conversion of 1 ug genomic DNA was performed using the EZ-96 DNA Methylation Kit (Deep Well Format) (Zymo Research; Irvine, CA, USA) according to the manufacturer’s instructions (www.zymoresearch.com). Bisulfite conversion efficiency was determined by PCR amplification of the converted DNA before proceeding with methylation analyses on the Illumina platform using Zymo Research’s Universal Methylated Human DNA Standard and Control Primers.

##### Normalization and Quality Control

Probes included on the list of cross-reactive 450K probes as reported by Chen *et al* were removed prior to analysis[37]. The cross-reactive target had to match a minimum of 47 bases to be considered cross-reactive. This led to the removal of ∼28,000 probes. Genome studio background correction and BMIQ normalization were performed[38] and the wateRmelon R package was used to conduct QC filtering[39].

Samples were removed for the following reasons: 1. Failed bisulfite conversion, 2. Call rate <95%, 3. Sex mismatch using minfi, 4. Weak correlation between available genotypes and genotypes on 450K array, 5. Weak clustering according to sex in MDS plot, 6. PCA analysis identified them as an outlier (≥4SD from mean), 7. Failed sex check, 8. Sample pass rate <99%, 9. Only sample to pass on a chip. These filtering settings led to the removal of 68 samples in the AA group and 24 samples in the EA group. If samples were run in duplicate, the sample with the lowest missing rate was retained.

##### Surrogate Variable Analysis (SVA)

SVAs were generated using the package SVA in R and protecting mtDNA-CN[36].

##### Control Probe Principal Components in ARIC European Americans

The control probe principal components are based on 42 measures, which are transformed from control probes and out-of-band probes in the 450K data[40].

#### Statistical Analysis

All statistical analyses were performed using R (version 3.3.3).

##### Linear Mixed Model – Association between mtDNA-CN and nuclear DNA methylation

Linear-mixed-effects regression analysis was performed to determine the association between mtDNA-CN and nuclear DNA methylation at specific CpGs (Supplemental Table S2).

**ARIC AA:** Methylation ∼ MtDNA-CN + Age + Sex + Site + Visit + Chip Position + Plate + CD8 Count + CD4 Count + B-Cell Count + Monocyte Count + Granulocyte Count + Smoking Status + First 10 Surrogate Variables + Chip (as random effect).

**ARIC EA:** Same model as ARIC AA but further inclusion of Project (Brain MRI or Omics) as well as the first 10 PCs derived from methylation microarray control probes and the composition of natural killer (NK) cells.

Cell types were imputed using the method of Houseman *et al*.[41]. All correlations were performed using the Pearson method.

Global methylation distributions were assessed by a chi-square test to compare observed to expected site-specific methylation.

##### Discovery Meta-Analysis

A meta-analysis was performed to combine the results from the individual ARIC AA and EA analyses (Supplemental Table S2). This analysis was done using the standard error scheme implemented in Metal[42]. CpGs had to have a *P*-value cutoff of *P*<0.05 in ARIC AA and EA analyses to be included in the meta-analysis. Associations that met genome-wide significance were included in subsequent analyses (*P*=5.0×10^-8^). 100 meta-analysis permutations were also performed (Permuted *P*=3.94×10^-8^).

##### Residual Bootstrapping

Residual bootstrapping was used to determine the most appropriate genome-wide significance cutoff in ARIC EA and AA cohorts (AA: *P*<6.22×10^-8^, EA: *P*<3.03×10^-7^) (see supplemental methods). Additionally, the qq plots show minimal inflation in ARIC AA, EA and meta-analysis (Supplemental Figure S1).

Significant CpGs with high correlation (R^2^≥0.6) were identified as non-independent and the CpGs with the more significant *P*-value was retained. Highly correlated CpGs were consistent between AA and EA results, specifically these CpGs were cg21051031 and cg03964851 (R^2^: AA=0.62, EA=0.63) and cg06809544 and cg13393978 (R^2^: AA=0.65, EA=0.70).

### Validation Cohorts

#### The Cardiovascular Health Study (CHS)

The CHS is a population-based cohort study of risk factors for coronary heart disease and stroke in adults ≥65 years conducted across four field centers[43]. The original predominantly European ancestry cohort of 5,201 persons was recruited in 1989-1990 from random samples of the Medicare eligibility lists; subsequently, an additional predominantly African-American cohort of 687 persons was enrolled in 1992-1993 for a total sample of 5,888. The validation cohort includes 239 AA participants and 294 EA participants from CHS with mtDNA-CN and 450K methylation derived from the same visit (Supplemental Table S1).

##### mtDNA-CN Estimation using Quantitative PCR

mtDNA copy number (mtDNA-CN) was determined utilizing a multiplexed real time quantitative polymerase chain reaction (qPCR) assay with ABI TaqMan chemistry (Applied Biosystems) as previously described[7] (see supplemental methods). The final measure of mtDNA-CN is represented as the standardized residuals from a race-stratified mixed linear regression adjusting for age, sex, and sample collection site.

##### Methylation Analysis

Methylation measurements were performed at the Institute for Translational Genomics and Population Sciences at the Harbor-UCLA Medical Center Institute for Translational Genomics and Population Sciences (Los Angeles, CA). DNA was extracted from Buffy coat fractions and subsequently underwent bisulfite conversion using the EZ DNA Methylation kit (Zymo Research, Irvine, CA). Methylation was then assayed using the Infinium HumanMethylation450 BeadChip (Illumina Inc, San Diego, CA) (see supplemental methods).

##### Regression Analysis

CHS was analyzed using linear regression with methylation beta values as the dependent variable and mtDNA-CN as the independent variable. Analyses were adjusted for age, sex, batch, measured white blood cell count and estimated cell type counts.

#### The Framingham Heart Study (FHS)

FHS is a prospective study of individuals from Framingham, Massachusetts[44]. The validation cohort includes 1,995 EA participants from FHS with mtDNA-CN and 450K methylation derived from the same visit (Supplemental Table S1).

##### mtDNA-CN Estimation from Whole Genome Sequencing

mtDNA copy number (mtDNA-CN) was determined from whole genome sequencing data. Cohort-specific mtDNA-CN residuals were obtained by regressing mtDNA-CN on age, sex, and WBC counts. Mitochondrial DNA copy number was estimated by applying the fastMitoCalc software[45] to harmonized build 37 mappings of TOPMed deep whole genome sequencing data (freeze 5). The estimated mitochondrial copy number is twice the ratio of average mitochondrial sequencing depth to average autosomal sequencing depth. We applied inverse normal transformation to mtDNA-CN residuals.

##### Methylation Analysis

DNA extraction, methylation quantification (450k-BeadChip), and QC were detailed previously[46]. We obtained lab-specific and cohort-specific DNA methylation residuals by regressing methylation beta values on age, sex, batch effects (plate, col, row), and WBC counts. We applied inverse normal transformation to DNA methylation residuals.

##### Regression Analysis

A linear mixed model was applied with inverse normal transformed DNA methylation residuals as the dependent variable and inverse normal transformed mtDNA-CN residuals as the independent variable, accounting for family structure.

#### Validation and all-cohort meta-analyses

A meta-analysis was performed of all validation cohorts (FHS EA, CHS EA, CHS AA). We also performed an all-cohort meta-analysis (ARIC AA, ARIC EA, FHS EA, CHS EA, CHS AA). Both meta-analyses were performed using the standard error scheme implemented in Metal[42]. Cohort specific data is available through the respective coordinating centers.

### Mendelian Randomization

#### meQTL Analysis

meQTLs were identified using MatrixEQTL[47]. Imputed genotypes which were previously derived from ARIC for the relevant participants as well as normalized residuals from our 450K methylation dataset were used in regression analysis (see supplemental methods). Metasoft[48] was used for meta-analysis; in addition to the fixed effects (FE) model, a random effects (RE) and Han and Eskin’s Random Effects model (RE2) were also used and yielded very similar results (Supplemental Table S9).

#### Mendelian Randomization Methods

Independent meQTLs were used for MR. Independence was defined by including SNPs in the same linear model. MR with mtDNA-CN as the outcome and methylation as the exposure was undertaken. meQTLs served as the known relationship of genotype on exposure (methylation) and the results of the linear model, lm(mtDNA∼meQTL SNP) were calculated. Power for the MR was calculated using the YZ association function in mRnd[49].

### Phenotype Analysis

We compared methylation at the six validated CpGs to phenotypes that are known to be associated with mtDNA-CN. Phenotypes included prevalent diseases (CHD, CVD) as well as incident diseases (CHD, CVD, Mortality). The analysis was performed as follows for each cohort:

A) Prevalent diseases (CHD, CVD): glm(PRVCVD ∼ resids(methyl) + AGE + SEX + CENTER + RACE, family=binomial(logit))
B) Incident diseases (CHD, CVD, Mortality): coxph(Surv(STime,dead) ∼ resids(methyl) + AGE + SEX + CENTER + RACE))

Where resids(methyl) represents methylation adjusted for all relevant covariates from the EWAS. The event adjudication process in ARIC, CHS and FHS consisted of expert committee review of hospital records, telephone interviews, and death certificates (see supplemental methods).

Results from each of the five individual cohorts were meta-analyzed across cohorts using an inverse weighted standard error method[42] to derive an overall phenotype association for each CpG of interest.

### CRISPR-Cas9 Knockout of TFAM

#### Generation of *TFAM* Knockout

The stable *TFAM* CRISPR-Cas9 knockout was generated in HEK293T cells using the Origene *TFAM* – Human Gene Knockout Kit via CRISPR (catalog number: KN215488) following the manufacturer’s protocol. The following sgRNA guide sequence was used to generate the stable *TFAM* knockout lines: GCGTTTCTCCGAAGCATGTG. Lipofection was conducted using Turbofectin 8.0 (catalog number: TF81001). Puromycin was used for selection at a concentration of 1.5 µg/mL. Fluorescence-activated cell sorting (FACS) was used for single cell sorting and clonal expansion. HEK293T cells were grown in DMEM containing 10% FBS and 1% penicillin-streptomycin at 37°C and 5% CO2. Sequencing primers used to confirm the *TFAM* knockout and proper insertion of the Donor plasmid are as follows: *TFAM*_Left_Forward_Primer_2: AGCGACTGTGGACAACTAGC, GFP_Reverse_Primer_2: TCATCTTGTTGGTCATGCGG, Puro-Forward_Primer_1: CACAACCTCCCCTTCTACGAG, *TFAM*_Right_Reverse_Primer_1: CCCCAAACTCCTTACCTGGG. DNA/RNA/protein isolation, mtDNA-CN estimation, qPCR to determine *TFAM* DNA quantity and *TFAM* expression as well as Western blotting were performed (see supplemental methods).

#### Methylation Analysis of *TFAM* Knockout Lines

*TFAM* KO cell lines were hybridized to the Illumina Infinium EPIC BeadChip at The University of Texas Health Science Center at Houston (UTHealth). Bisulfite conversion efficiency was reviewed in the laboratory using the Bead Array Controls Reporter (BACR) tool, and Illumina chemistry (sample independent controls) performed within acceptable specifications.

All samples passed with detected CpG (0.01) >97%.

EPIC BeadChip analysis was performed using the minfi package[50]. Data was normalized using Functional Normalization[40] and differential methylation was calculated using the dmpFinder function in minfi (Supplemental Table S4).

In the cases where the CpG from the 450k array was not represented on the EPIC array a CpG surrogate was chosen if there was a nearby CpG within 1000 bp upstream or downstream of the original CpG that was highly correlated with the original CpG (R^2^ ≥0.6) and associated with mtDNA-CN in the ARIC analysis (*P*<5×10^-8^).

#### RNA sequencing of *TFAM* Knockout Lines

##### RNA Preparation

RNA quantification was performed using the Qubit RNA BR Assay (Invitrogen #Q10211) and Qubit 2.0 Fluorometer. The Agilent BioAnalyzer was used for quality control of the RNA prior to library creation, with a minimum RIN of 8.5. Samples were diluted to 300 ng/uL in 12 uL molecular biology grade water, and then submitted to the Genetic Resources Core Facility for RNA sequencing. RNA-seq resulted in expected clustering of knockout and control lines (Supplemental Figure S10).

##### Library Preparation and Sequencing

Illumina’s TruSeq Stranded Total RNA kit protocol was used to generate libraries (see supplemental methods).

##### Primary Analysis

Illumina HiSeq reads were processed through Illumina’s Real-Time Analysis (RTA) software generating base calls and corresponding base call quality scores. CIDRSeqSuite 7.1.0 was used to convert compressed bcl files into compressed fastq files.

##### Secondary Analysis

Each independent cell-line was sequenced twice. RNA sequencing fastq files were pseudoaligned to Genome Reference Consortium Human Build 37 (GRCh37) using Kallisto[51]. 100 bootstraps were performed using Kallisto. The R package Sleuth was used for RNA sequencing analysis[52] (Supplemental Table S5). Lane was included as a covariate in the Sleuth model. Differentially expressed genes were defined as those with a *P*<0.05.

#### Integrated analysis of *TFAM* knockout methylation and expression

The linear-gwis method in FAST (genotype mode) was used to collapse *TFAM* KO methylation data into one gene level *P*-value per gene[53]. These gene-level methylation results were combined with gene-level gene expression results for the same gene using the Fisher *P*-value combination method to generate an integrated gene level Methylation/RNA sequencing *P*-value.

### GO/KEGG Analysis

Each CpG was annotated with the nearest gene as defined by the closest gene which harbors the CpG within 1,500 bp of the transcriptional start site and extending to the polyA signal. A bias exists when performing gene set analysis for genome-wide methylation data that occurs due to the differing numbers of CpG sites profiled for each gene[54]. Due to this, we used gometh for GO and KEGG analysis since it is based off of the goseq method which accounts for this bias[55]. We analyzed our individual ARIC/*TFAM* datasets as well as our *TFAM* integrated (meth/expression) dataset. We also combined GO/KEGG results for ARIC, *TFAM* methylation and *TFAM* RNA sequencing using the Fisher *P*-value combination method to generate an overall combined *P*-value for each term. Final *P*-value cutoffs used for each analysis were as follows: ARIC Discovery Meta-Analysis (300 CpGs, *P*=5.24×10^-12^), *TFAM* Methylation (300 CpGs, *P*=4.41×10^-4^), *TFAM* Expression (169 genes, *P*=4.30×10^-4^), *TFAM* Integrated (Methylation/Expression) (188 genes, *P*=8.77×10^-6^).

## LIST OF ABBREVIATIONS

AA: African American
ARIC: Atherosclerosis Risk in Communities
CHD: Coronary Heart Disease
CHS: Cardiovascular Health Study
CVD: Cardiovascular disease
EA: European American
FHS: Framingham Heart Study
meQTLs: methylation quantitative trait loci
MR: Mendelian Randomization
mtDNA: mitochondrial DNA
mtDNA-CN: Mitochondrial DNA copy number
nDNA: nuclear DNA
qPCR: quantitative polymerase chain reaction
SNPs: single nucleotide polymorphisms
SVA: Surrogate Variable Analysis
TOPMed: Trans-Omics in Precision Medicine
WGS: Whole genome sequencing

## DECLARATIONS

### ETHICS APPROVAL AND CONSENT TO PARTICIPATE

The Atherosclerosis Risk in Communities (ARIC) study, Cardiovascular Health Study (CHS) and Framingham Heart Study (FHS) have been approved by the Institutional Review Board (IRB) at each participating institution. All participants provided written informed consent.

The ARIC study design and methods were approved by four different IRBs at each of the collaborating medical institutions: University of Mississippi Medical Center Institutional Review Board (Jackson Field Center); Wake Forest University Health Sciences Institutional Review Board (Forsyth County Field Center); University of Minnesota Institutional Review Board (Minnesota Field Center); and Johns Hopkins University School of Public Health Institutional Review Board (Washington County Field Center).

FHS is approved by the IRB at Boston University Medical Center. CHS recruited participants from Medicare lists at four sites and IRBs at each site were involved in human subjects approval.

### CONSENT FOR PUBLICATION

Not applicable.

### AVAILABILITY OF DATA AND MATERIALS

All raw and processed sequencing data generated and analyzed during the current study are available in the NCBI Gene Expression Omnibus (GEO; http://www.ncbi.nlm.nih.gov/geo/) under accession numbers GSE133994 and GSE134048.

### COMPETING INTERESTS

The authors declare that they have no competing interests.

### FUNDING

This research was supported by grant R01HL131573 from the US National Institutes of Health. Castellani was supported by a CIHR Postdoctoral Fellowship. Cohort funding information can be found in the cohort specific acknowledgement sections below.

### AUTHOR CONTRIBUTIONS

C.A.C., E.G., N.P., B.O., J.C., and D.E.A. conceived and designed the experiments. C.A.C., R.J.L., C.E.N., J.A.S., J.A.L., J.A.B., T.M.B., M.L.G., M.F., J.S.F., J.B., J.S.P, A.T., B.O., E.G., N.P., K.D.T., P.W., C.L., E.B., and D.E.A. acquired, analyzed and interpreted the data. C.A.C. and D.E.A. were responsible for the drafting of the manuscript. C.A.C., R.J.L., J.S.F., C.L., A.T., M.F., B.O., J.A.B., J.S.P, T.M.B. and D.E.A. critically revised the manuscript for important intellectual content. C.A.C., R.J.L., J.A.L., J.A.B., C.L., E.G., N.P., and D.E.A. performed statistical analysis. J.C., E.G., E.B. and D.E.A. obtained funding. C.E.N., J.A.S., M.L.G., J.B. provided administrative, technical, or material support. This work was performed under the supervision of N.S., D.L., E.G., and D.E.A. All authors read and approved the final manuscript.

## ACKNOWLEDGEMENTS

Infinium Methylation EPIC BeadChip array hybridization was performed at the UTHealth Human Genetics Center, The University of Texas, Houston, TX. Illumina sequencing was conducted at the Genetic Resources Core Facility, Johns Hopkins Institute of Genetic Medicine, Baltimore, MD.

### ARIC Acknowledgements

The Atherosclerosis Risk in Communities study has been funded in whole or in part with Federal funds from the National Heart, Lung, and Blood Institute, National Institutes of Health, Department of Health and Human Services (contract numbers HHSN268201700001I, HHSN268201700002I, HHSN268201700003I, HHSN268201700004I and HHSN268201700005I). The authors thank the staff and participants of the ARIC study for their important contributions. Funding was also supported by 5RC2HL102419 and R01NS087541.

### CHS Acknowledgements

Infrastructure for the CHARGE Consortium is supported in part by the National Heart, Lung, and Blood Institute grant R01HL105756. The CHS research was supported by NHLBI contracts HHSN268201200036C, HHSN268200800007C, HHSN268201800001C, N01HC55222, N01HC85079, N01HC85080, N01HC85081, N01HC85082, N01HC85083, N01HC85086; and NHLBI grants U01HL080295, U01HL130114, K08HL116640, R01HL087652, R01HL092111, R01HL103612, R01HL103612, R01HL111089, R01HL116747 and R01HL120393 with additional contribution from the National Institute of Neurological Disorders and Stroke (NINDS). Additional support was provided through R01AG023629 from the National Institute on Aging (NIA), Merck Foundation / Society of Epidemiologic Research as well as Laughlin Family, Alpha Phi Foundation, and Locke Charitable Foundation. A full list of principal CHS investigators and institutions can be found at CHS-NHLBI.org. The provision of genotyping data was supported in part by the National Center for Advancing Translational Sciences, CTSI grant UL1TR000124, and the National Institute of Diabetes and Digestive and Kidney Disease Diabetes Research Center (DRC) grant DK063491 to the Southern California Diabetes Endocrinology Research Center. The content is solely the responsibility of the authors and does not necessarily represent the official views of the National Institutes of Health.

### FHS Acknowledgements

Whole genome sequencing (WGS) for the Trans-Omics in Precision Medicine (TOPMed) program was supported by the National Heart, Lung and Blood Institute (NHLBI). WGS for “NHLBI TOPMed: Whole Genome Sequencing and Related Phenotypes in the Framingham Heart Study” (phs000974.v1.p1) was performed at the Broad Institute of MIT and Harvard (HHSN268201500014C). Centralized read mapping and genotype calling, along with variant quality metrics and filtering were provided by the TOPMed Informatics Research Center (3R01HL-117626-02S1). Phenotype harmonization, data management, sample-identity QC, and general study coordination, were provided by the TOPMed Data Coordinating Center (3R01HL-120393-02S1). We gratefully acknowledge the studies and participants who provided biological samples and data for TOPMed.

This work is supported by National Institutes of Health (NIH) contract N01-HC-25195 and HHSN268201500001I and grant R01 HL092577, also supported by intramural funding of Dan Levy, National Heart, Lung, and Blood Institute (NHLBI) (for DNA methylation profiling), and Trans-Omics for Precision Medicine (TOPMed) sponsored by NHLBI/NIH. The Framingham Heart Study thanks the study participants and the multitude of investigators who over its 70 year history continue to contribute so much to further our knowledge of heart, lung, blood and sleep disorders and associated traits.

The views expressed in this manuscript are those of the authors and do not necessarily represent the views of the National Heart, Lung, and Blood Institute; the National Institutes of Health; or the U.S. Department of Health and Human Services.

## ADDITIONAL MATERIAL

**Supplementary Materials.pdf** →Supplementary Methods and Supplemental Figures:

**Figure S1.** QQ plots from ARIC association analyses.

**Figure S2.** Manhattan plot of association between nuclear DNA methylation and mtDNA-CN in ARIC (discovery cohort) stratified analysis.

**Figure S3.** Correlation between beta estimates for significant CpGs identified by ARIC EA and ARIC AA analyses (Discovery cohort)

**Figure S4.** Volcano Plots of results from ARIC association analyses.

**Figure S5.** Cohort correlations between beta estimates for genome-wide significant CpGs in discovery cohort (ARIC) and beta estimates from CHS/FHS validation cohorts.

**Figure S6.** Individual cohort results for 6 validated CpGs.

**Figure S7.** qPCR DNA Copy Number results for TFAM gene in negative control and TFAM knockout lines.

**Figure S8.** ARIC identified CpGs with methylation differences between EPIC Negative Control (NC) lines and TFAM knockout (CRISPR) cell lines.

**Figure S9.** Global methylation patterns in negative Control (NC) and CRISPR-TFAM knockout (CR) cell lines.

**Figure S10.** TFAM knockout RNA sequencing clustering heatmap of TFAM knockout and control cell lines.

**Supplemental_Table_S1.pdf** → Sample characteristics of discovery and validation cohorts.

**Supplemental_Table_S2.xlsx** →ARIC summary statistics. A. Meta-Analysis, B. AA, C. EA.

**Supplemental_Table_S3.xlsx**→A. Results for 34 independent ARIC Discovery Meta-Analysis identified mtDNA-CN associated CpGs across all studied cohorts and Validation Meta-Analysis/All Cohort Meta-Analysis. Validation meta-analysis included CHS AA, CHS EA and FHS EA cohorts (P<0.05 and same direction, bolded cells). All cohort meta-analysis (ARIC AA, ARIC EA, CHS AA, CHS EA and FHS EA) identified 6 validated CpGs (P<5×10-8, shaded cells). B. Results for 23 independent ARIC AA identified CpGs across all studied cohorts. C. Results for 15 independent ARIC EA identified CpGs across all studied cohorts.

**Supplemental_Table_S4.xlsx**→TFAM Methylation analysis summary statistics.

**Supplemental_Table_S5.xlsx**→TFAM Expression analysis summary statistics.

**Supplemental_Table_S6.pdf** →Neuroactive-ligand receptor interaction genes as identified by KEGG analysis in each approach, number of genes used in each analysis and neuroactive ligand enrichment P-value are included in brackets.

**Supplemental_Table_S7.pdf**→Methylation Status of Validated CpGs in TFAM KO cell lines (N=6). Bolded entries indicate differential expression P<0.05.

**Supplemental_Table_S8.pdf**→ Differentially expressed genes (*P*<0.05) within 1 Mb of differentially methylated CpGs in *TFAM* knockout cell lines. Shading indicates most differentially expressed gene for each CpG.

**Supplemental_Table_S9.pdf**→**Results of Mendelian Randomization. A.** Results for association between ARIC EA and AA derived independent *cis* meQTLs and mtDNA-CN. **B.** Results for association between ARIC meta-analysis derived independent *cis* meQTLs and mtDNA-CN (fixed effects model).

**Supplemental_Table_S10.xlsx** → Phenotype analysis: Cohort specific results and meta-analysis of all cohorts.

