## Supplemental Methods and Figures for "Mitochondrial DNA Copy Number (mtDNA-CN) Can Influence Mortality and Cardiovascular Disease via Methylation of Nuclear DNA CpGs"

### Supplemental Materials

### Supplementary Methods

##### *Ethics*

The Atherosclerosis Risk in Communities (ARIC) study, Cardiovascular Health Study (CHS) and Framingham Heart Study (FHS) have been approved by the Institutional Review Board (IRB) at each participating institution. All participants provided written informed consent.

The ARIC study design and methods were approved by four different IRBs at each of the collaborating medical institutions: University of Mississippi Medical Center Institutional Review Board (Jackson Field Center); Wake Forest University Health Sciences Institutional Review Board (Forsyth County Field Center); University of Minnesota Institutional Review Board (Minnesota Field Center); and Johns Hopkins University School of Public Health Institutional Review Board (Washington County Field Center).

FHS is approved by the IRB at Boston University Medical Center. CHS recruited participants from Medicare lists at four sites and IRBs at each site were involved in human subjects approval.

**Statistical Analysis**

*Residual Bootstrapping*

The steps taken were as follows: 1) Residuals were derived from the full model, 2) Fitted values were derived from the null model (model without mtDNA-CN as independent factor), 3) The residuals from Step 1 were resampled and added to the fits from Step 2, 4) Each resulting matrix from Step 3 was run as pseudonull input in the formula lme(pseudonull~CN+covariates) to refit the full model and obtain null statistics, 4a) The most extreme P-value was pulled from each iteration, 4b) The resulting 100 most extreme P-values were ranked from least to most significant and the 95th value was chosen to be the 'genome-wide significance level' for the corresponding cohort.

***Validation Cohorts***

**The Cardiovascular Health Study (CHS)**

*mtDNA-CN Estimation using Quantitative PCR*

Briefly, each well consisted of a VIC-labeled, primer limited assay specific to a mitochondrial target (*MT-ND1*), and a FAM-labeled assay specific to a region of the nuclear genome selected for being non-repetitive (*RPPH1*). Each sample was run in triplicate on a 384 well plate in a 10 $\mu$L reaction containing 20 ng of DNA. The cycle threshold (Ct) value was determined from the amplification curve for each target by the ABI Viia7 software. A ΔCt value was computed for each well as the difference between the Ct for the *RPPH1* target and the Ct for the *MT-ND1* target, as a measure of mtDNA copy number relative to nuclear DNA copy number. For samples with a standard deviation of ΔCt for the three replicates >0.5, an outlier replicate was identified and excluded. If the ΔCt standard deviation remained >0.5 after exclusion, the sample was completely excluded from future analyses. Replicates with Ct values for *MT-ND1* >28, Ct values for *RPPH1* >5 standard deviations from the mean, or ΔCt values >3 standard deviations from the mean of the plate were removed. Additionally, due to an observed linear increase in ΔCt value by the order in which the replicate was pipetted onto the plate, a linear regression was used to correct for pipetting order. Plate effects are controlled for by performing a linear regression whereby the plate a sample is run on is treated as a random effect.

*Methylation Analysis*

Quality control was performed in in the minfi R package(Aryee et al. 2014) (version 1.12.0, http://www.bioconductor.org/packages/release/bioc/html/minfi.html). Samples with low median intensities of below 10.5 (log2) across the methylated and unmethylated channels, samples with a proportion of probes falling detection of greater than 0.5%, samples with QC probes falling greater than three standard deviations from the mean, sex-check mismatches, failed concordance with prior genotyping or > 0.5% of probes with a detection *P*-value > 0.01 were removed. Probes with >1% of values below detection were removed. In total, 11 samples were removed for sample QC resulting in a sample of 323 European-ancestry and 326 African-American samples. Methylation values were normalized using the SWAN quantile normalization method(Maksimovic et al. 2012). Since white blood cell proportions were not directly measured in CHS they were estimated from the methylation data using the Houseman method(Houseman et al. 2014).

***Mendelian Randomization***

**meQTL Analysis**

Haplotype phasing was performed using ShapeIt(Delaneau et al. 2013) and imputation was performed using IMPUTE2(Howie et al. 2012). SNPs were filtered for allele frequency >0.05, and imputation quality >0.4. Genotypes were imputed to the 1000G reference panel (Phase I, version 3). The same covariates used for the ARIC EWAS analysis were used to call meQTLs as well as the addition of genotyping PCs (4 for EA, 10 for AA). Only meQTLs which had an individual cohort *P* value >0.05 were included in the meta-analysis.

A linear model was used for MatrixEQTL and a cis meQTL was defined as having a distance less than 100 kb. Only cis meQTLs derived from the six CpGs of interest and which met a cohort-specific permuted *P*-value cutoff (Permuted *P:* EA=$7.84 \times 10^{-4}$, AA=$9.12 \times 10^{-4}$) or a permuted meta-analysis *P*-value cutoff (Permuted *P*, fixed effects (FE) model: $3.97 \times 10^{-5}$) were retained for use in Mendelian randomization.

***Phenotype Analysis***

Adjudicated events between visit 1 and the baseline visit for this study were considered prevalent events. Analyses of prevalent and incident events in CHS were adjusted for age, sex, clinic site and batch.

In ARIC, prevalent coronary heart disease (CHD) was defined as history of myocardial infarction (MI) or cardiac procedures (heart or arterial surgery, coronary bypass, or angioplasty). Cardiovascular disease (CVD) was defined as either CHD or stroke. Prevalent stroke was defined as stroke at baseline. For all phenotypes, prevalent disease was a combination of self-report at visit 1 plus adjudicated events between visit 1 and the baseline visit. Incident CHD was defined as the first incident MI or death owing to CHD. Incident stroke was defined as the first nonfatal stroke or death owing to stroke. In ARIC, the mean follow-up time was 20.6 years in the EA cohort and 18.1 years in the AA cohort. Follow-up for incident events was administratively censored at December 31, 2016.

CHS and FHS followed similar phenotype definitions as ARIC. For FHS, the mean follow-up time was 6.0 years and individuals were removed if follow-up years equaled 0, FHS events were adjudicated through 12/2016. In CHS, prevalent CVD/CHD was excluded during sampling and events were adjudicated through June 30, 2015. The follow up time for incident events from the time of methylation measurement was 23 years.

***CRISPR-Cas9 Knockout of TFAM***

*DNA Isolation*

DNA extraction was performed on harvested HEK293T *TFAM* knockout cells using the AllPrep DNA/RNA Mini Kit (Qiagen #80204) following the manufacturer's protocol (passage 32 and 37). DNA was eluted in 100 $\mu$L ultrapure water. DNA was quantified using a Nanodrop 1000. Low purity samples were subjected to ethanol precipitation.

*RNA Isolation*

Total RNA was extracted from confluent T75 culture flasks of *TFAM* CRISPR Negative Control and KO cell lines using the AllPrep DNA/RNA/Protein Kit (Qiagen #80004) (passage 45). RNA was extracted using the provided kit manual/instructions for RNA extraction, except all microcentrifuge spins were performed at 10,000 x g. RNA was eluted twice in 50 uL molecular biology grade water and stored in a -80C freezer.

*mtDNA-CN Estimation on TFAM Knockout Cell Lines*

qPCR was used to measure mtDNA-CN as described above for CHS in section “mtDNA-CN Estimation using Quantitative PCR”.

*qPCR to determine TFAM* DNA quantity

20 ng of total DNA from each cell line was used as input for a 10 μL volume reaction. *TFAM* DNA were amplified using TaqMan probe Hs00273372_s1 (20x, FAM-labeled, Applied Biosystems #4331182). *GAPDH* DNA served as a housekeeping reference control and was amplified with probe Hs03929097_g1 (20x, VIC-labeled, Applied Biosystems #4448489). Both probes were multiplexed together and all qPCR reactions were conducted at 50$^{o}$ C for 2 min, 95$^{o}$C for 10 min, and then 40 cycles of 95$^{o}$C for 15 sec and 60$^{o}$C for 1 min. DNA quantity was determined using double delta cycle threshold using *GAPDH* as the reference and displayed a 66% reduction of the *TFAM* gene.

*TFAM Expression Assay*

cDNA synthesis was performed with the SuperScript III First-Strand Synthesis System for RT-PCR (ThermoFisher #18080-051) following the manufacturer's protocol. 1.5 μg of total RNA from each cell line was used as input and primed with 50 ng random hexamers using the appropriate incubation conditions from the manufacture’s protocol. Following completed cDNA synthesis, samples were quantified using the Qubit ssDNA assay kit (Invitrogen #Q10212) and Qubit 2.0 Fluorometer. Synthesized cDNA was then diluted to 10 ng/μL using ultrapure water and stored in -20$^{o}$C.

*qPCR to determine TFAM* gene expression for *TFAM* KO

20 ng of synthesized cDNA from each cell line was used as input for a 10 μL volume reaction. *TFAM* cDNA were amplified using TaqMan probe Hs00273372_s1 (20x, FAM-labeled, Applied Biosystems #4331182). *GAPDH* cDNA served as a housekeeping reference control and was amplified with probe Hs03929097_g1 (20x, VIC-labeled, Applied Biosystems #4448489). Both probes were multiplexed together and all qPCR reactions were conducted at 50$^{o}$ C for 2 min, 95$^{o}$C for 10 min, and then 40 cycles of 95$^{o}$C for 15 sec and 60$^{o}$C for 1 min. Expression fold change was determined using double delta cycle threshold using *GAPDH* as the housekeeping reference control.

*Total Protein Extraction*

Total protein lysates from HEK293T *TFAM* CRISPR knockout cell lines were extracted using ice-cold radioimmunoprecipitation assay buffer (RIPA) buffer supplemented with Halt Protease and Phosphatase Inhibitor Cocktail (Thermo Scientific #78440) (passage 35). Protein concentrations were quantified using the Pierce BCA Protein Assay Kit (Thermo Scientific #23227) and lysates were stored at -80°C.

*Western Blotting*

Equal amounts of each lysate were diluted 1:1 with 2x Laemmli Sample Buffer (Bio-Rad #161-0737) supplemented with 5% β-mercaptoethanol. Samples were then heated at 95°C for 5 minutes to denature the proteins. 30 $\mu$g of each protein lysate was separated on a 12% polyacrylamide Mini-PROTEAN TGX Gel (Bio-Rad #456-1044) and then transferred to a PVDF membrane (Bio-Rad #1704156) using the Trans-Blot Turbo Transfer System. The membrane was blocked overnight at 4° C in Tris-Buffered Saline and Tween 20 (TBST) containing 5% nonfat milk with gentle shaking. After blocking, the membrane was incubated with rabbit anti-*TFAM* primary antibody diluted 1:40,000 in 5% milk (Abcam #ab176558) and rabbit anti- β-Tubulin primary antibody diluted 1:3,000 in 5% milk (Invitrogen #PA5-27552) for 1 hour at room temperature with gentle shaking. The membrane was washed 5-times with TBST after primary antibody incubation, then incubated with goat anti-rabbit secondary antibody conjugated with horseradish peroxidase (1:20,000 dilution, Abcam #ab97080) in the dark for 1 hour at room temperature with shaking. Signals were visualized by enhanced chemiluminescent substrate (SuperSignal West Pico PLUS, Thermo Scientific #34577) and photographed digitally using the ChemiDoc-It$^2$ Imager. ImageJ was used to quantify protein abundance (normalized to tubulin control).

**RNA sequencing of *TFAM* Knockout Lines**

*Library Preparation and Sequencing*

Specifically, total RNA is converted to cDNA and size selected to 150 to 200 bp in length with 3' or 5' overhangs. End repair is performed where 3' to 5' exonuclease activity of enzymes removes 3' overhangs and the polymerase activity fills in the 5' overhangs. An ‘A’ base is then added to the 3' end of the blunt phosphorylated DNA fragments which prepares the DNA fragments for ligation to the sequencing adapters, which have a single ‘T’ base overhang at their 3' end. Ligated fragments are subsequently size selected through purification using SPRI beads and undergo PCR amplification techniques to prepare the ‘libraries’. The BioAnalyzer is used for quality control of the libraries to ensure adequate concentration and appropriate fragment size. The resulting library insert size is 120-200 bp with a median size of 150 bp. Libraries were uniquely barcoded and pooled for sequencing. DNA sequencing was performed in duplicate on an Illumina® HiSeq 2500 instrument using standard protocols for paired end 150 bp sequencing. As per Illumina's recommendation, 3% PhiX was added to each lane as a control, and to assist the analysis software with any library diversity issues.

#### Supplemental Figures

**Figure S1. QQ plots from ARIC association analyses.** Observed *P*-values (black) versus expected *P*-values from a normal distribution (red). **A.** ARIC AA Cohort, **B.** ARIC EA Cohort. **C.** ARIC Meta-Analysis.

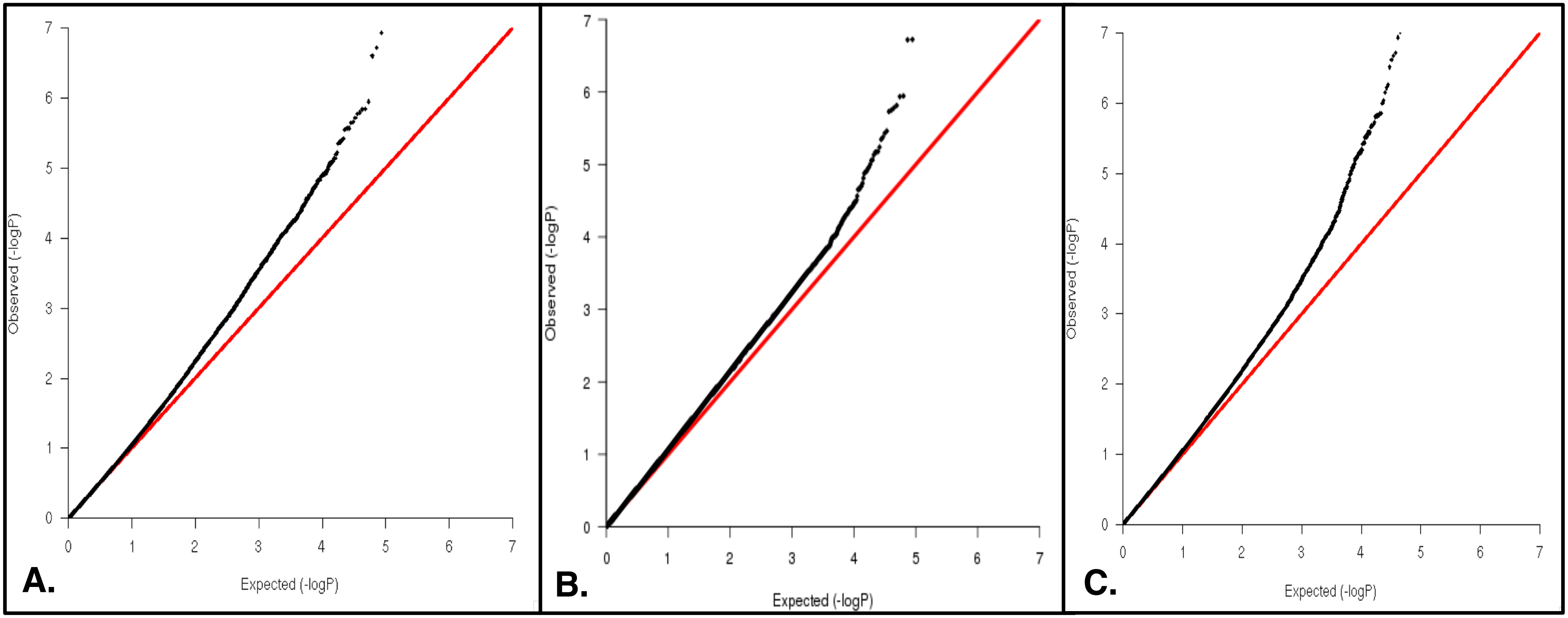

**Figure S2. Manhattan plot of association between nuclear DNA methylation and mtDNA-CN in ARIC (discovery cohort) stratified analysis. A.** ARIC AA, Blue dotted line represents cohort specific residual bootstrap cutoff ($P$=6.22x10$^{-8}$) **B.** ARIC EA, Blue dotted line represents cohort specific residual bootstrap cutoff ($P$=3.03x10$^{-7}$). Red dots highlight statistically significant CpGs.

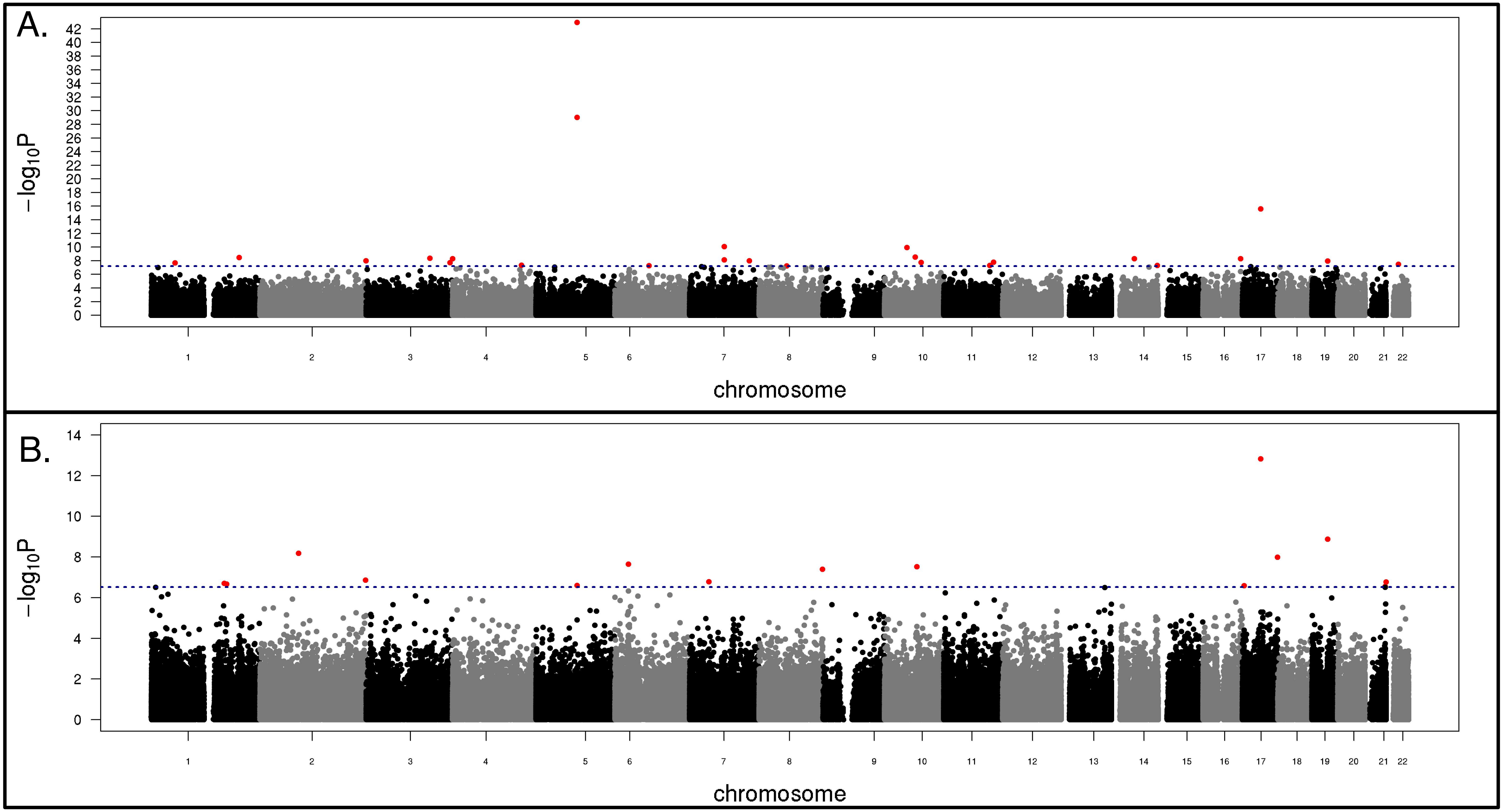

**Figure S3. Correlation between beta estimates for significant CpGs identified by ARIC EA and ARIC AA analyses (Discovery cohort).** Linear regression line shown in red ($R^2$=0.49), units are methylation betas. Green dots represent CpGs identified to be associated with mtDNA-CN in ARIC EA participants and purple dots represent CpGs identified to be associated with mtDNA-CN in ARIC AA participants.

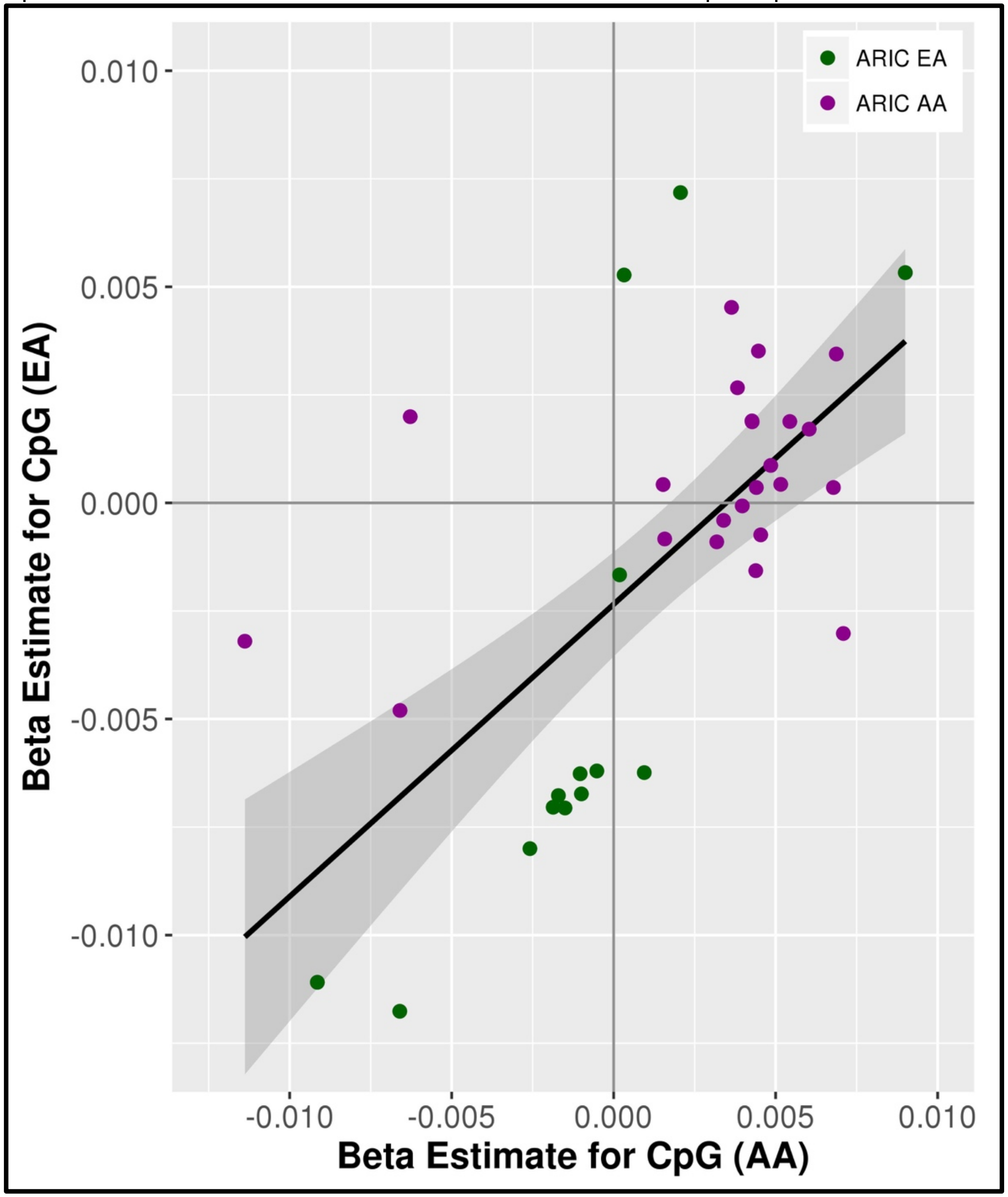

**Figure S4. Volcano Plots of results from ARIC association analyses. A.** ARIC AA Cohort **B.** ARIC EA Cohort **C.** ARIC Meta-analysis. Dark blue line represents *P*-value cut-off (AA $P=6.22\times10^{-8}$ , EA $P=3.03\times10^{-7}$, META $P=5.0\times10^{-8}$). Plots truncated at –log10(P)=10.

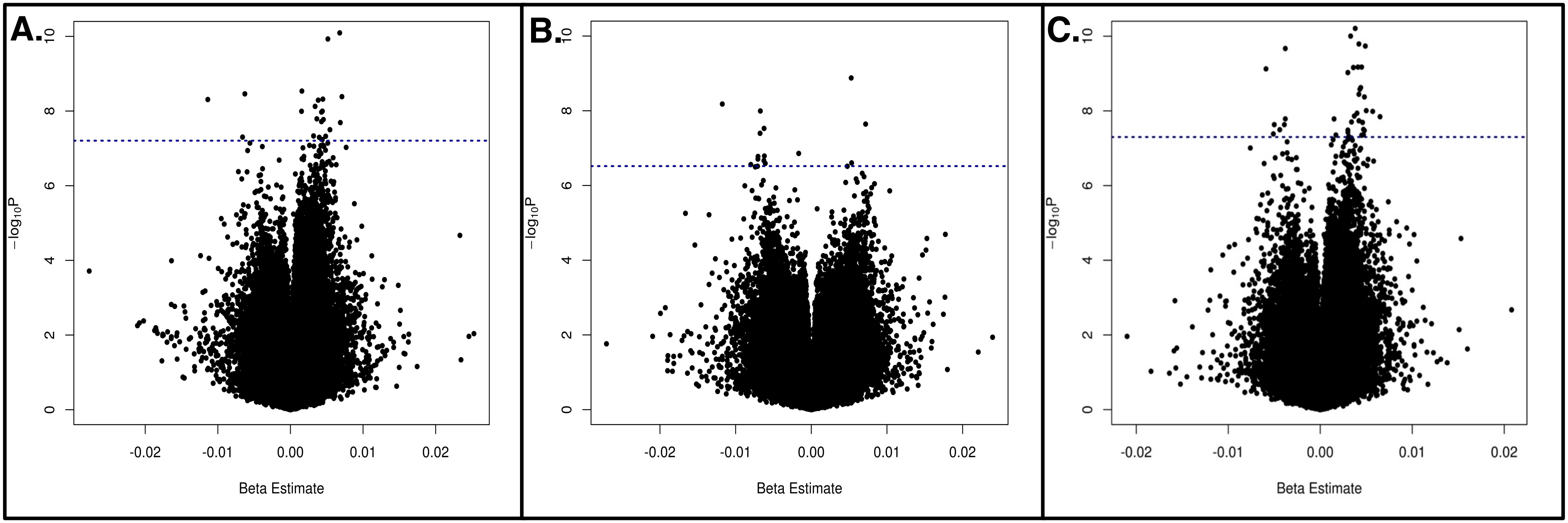

**Figure S5. Cohort correlations between beta estimates for genome-wide significant CpGs in discovery cohort (ARIC) and beta estimates from CHS/FHS validation cohorts. A.** ARIC Meta/CHS AA (*P* cutoff = $5.0 \times 10^{-8}$, r=0.26, Kendall tau=0.07) **B.** ARIC Meta/CHS EA (*P* cutoff = $5.0 \times 10^{-8}$, r=0.45, Kendall tau=0.10) **C.** ARIC Meta/FHS AA (*P* cutoff = $5.0 \times 10^{-8}$, r=0.58, Kendall tau=0.19) **D.** ARIC AA/CHS AA (*P* cutoff = $6.22 \times 10^{-8}$, r=0.55, Kendall tau=0.31) **E.** ARIC EA/CHS EA (*P* cutoff = $3.03 \times 10^{-7}$, r=0.01, Kendall tau=-0.07) **F.** ARIC EA/FHS EA (*P* cutoff = $3.03 \times 10^{-7}$, r=0.45, Kendall tau=0.27). Units are methylation betas.

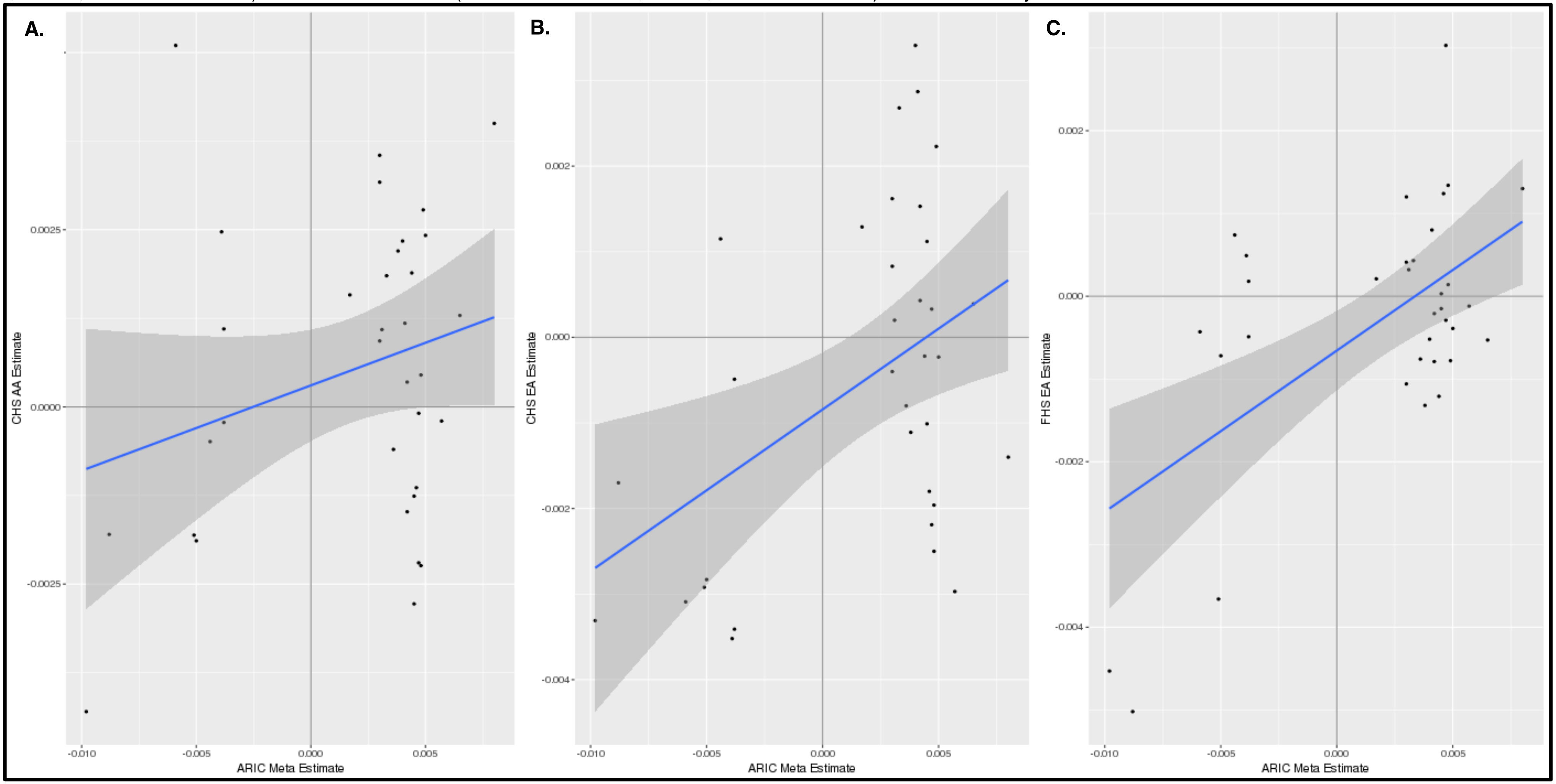

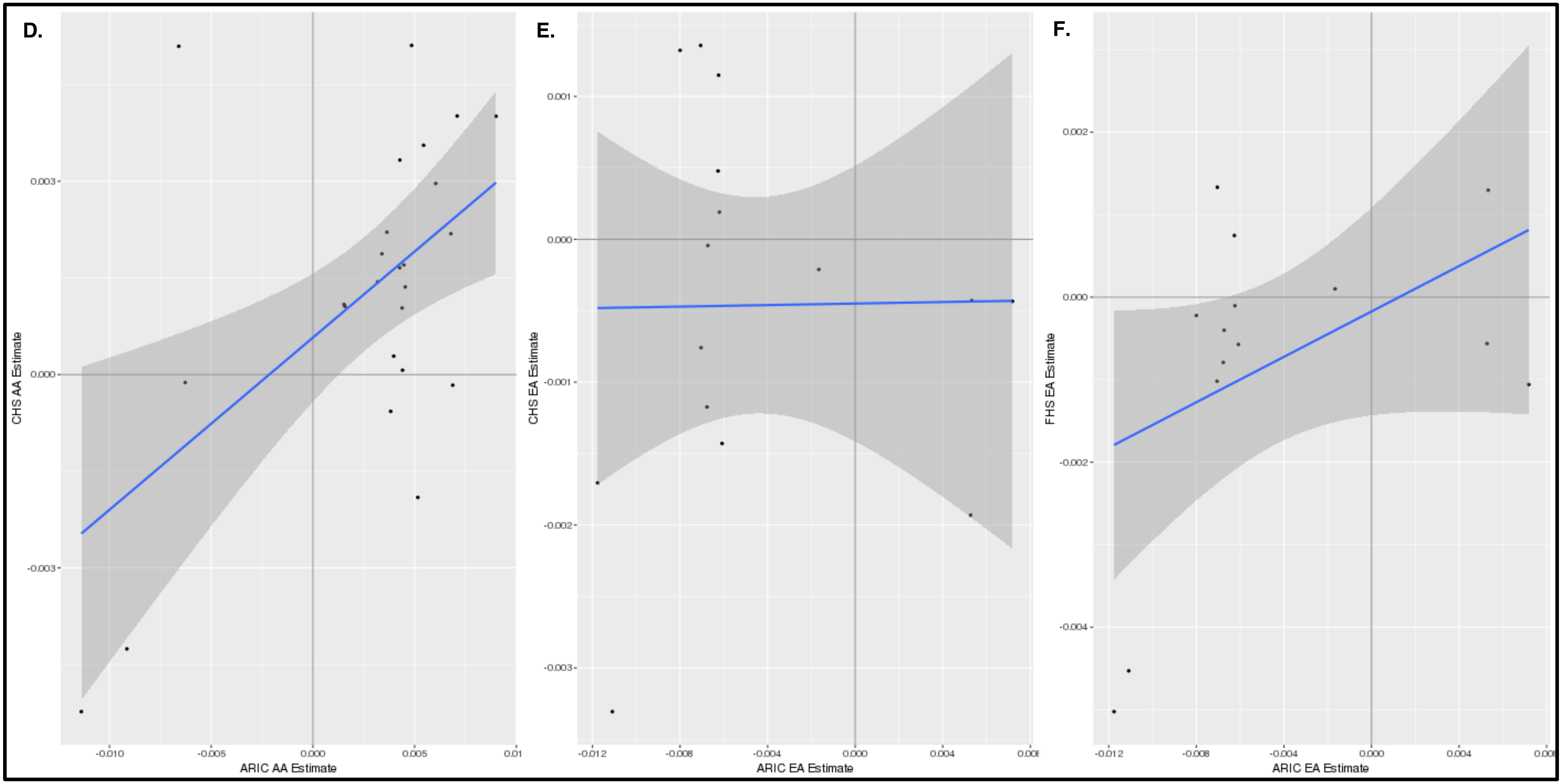
D.
CHS AA Estimate
ARIC AA Estimate
E.
CHS EA Estimate
ARIC EA Estimate
F.
FHS EA Estimate
ARIC EA Estimate

**Figure S6. Individual cohort results for 6 validated CpGs.**

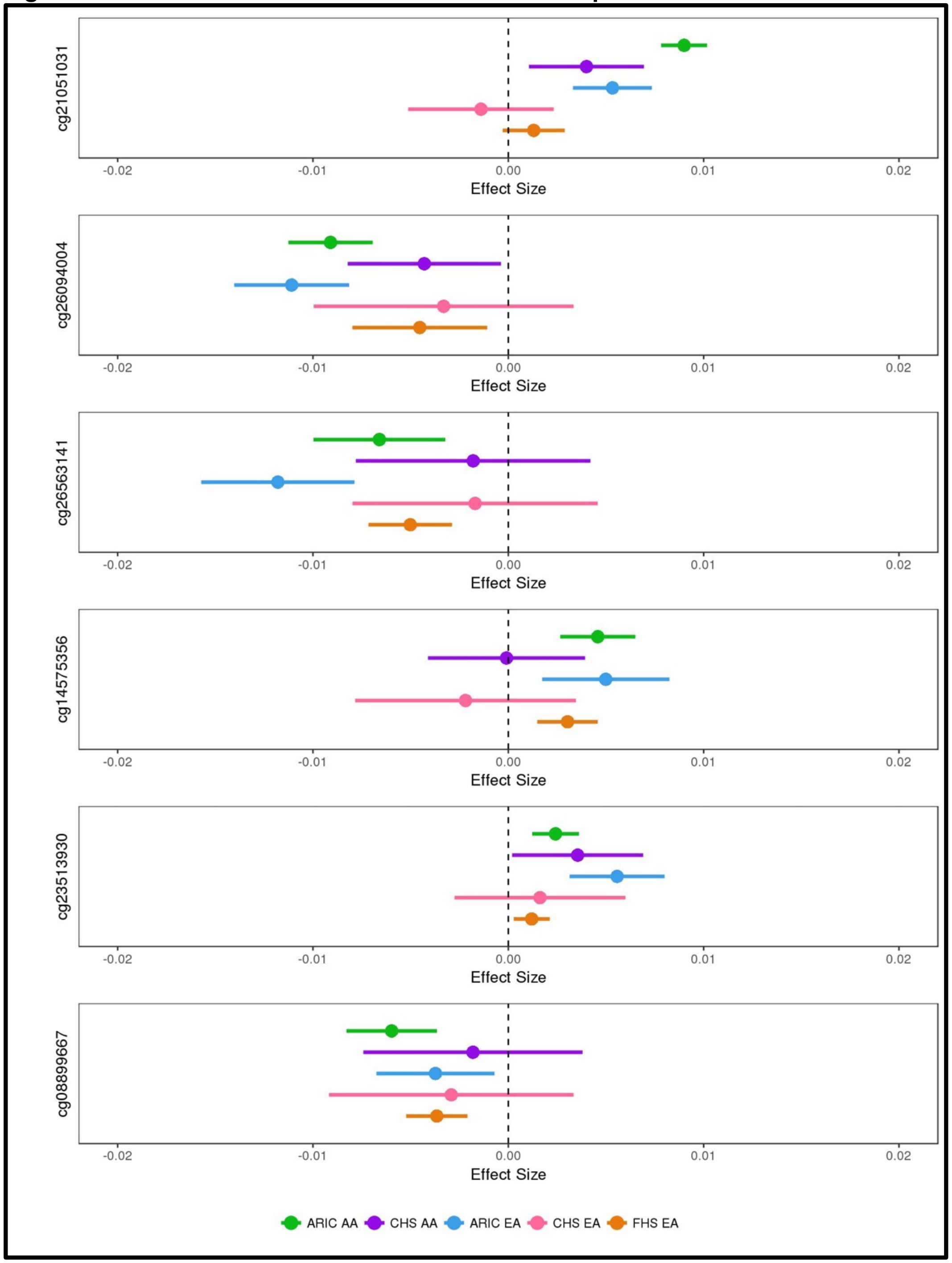

**Figure S7. qPCR DNA Copy Number results for *TFAM* gene in negative control and TFAM knockout lines.**
Control is *GAPDH*, △△Ct (p32)=1.57, △△Ct (p37)=1.56.

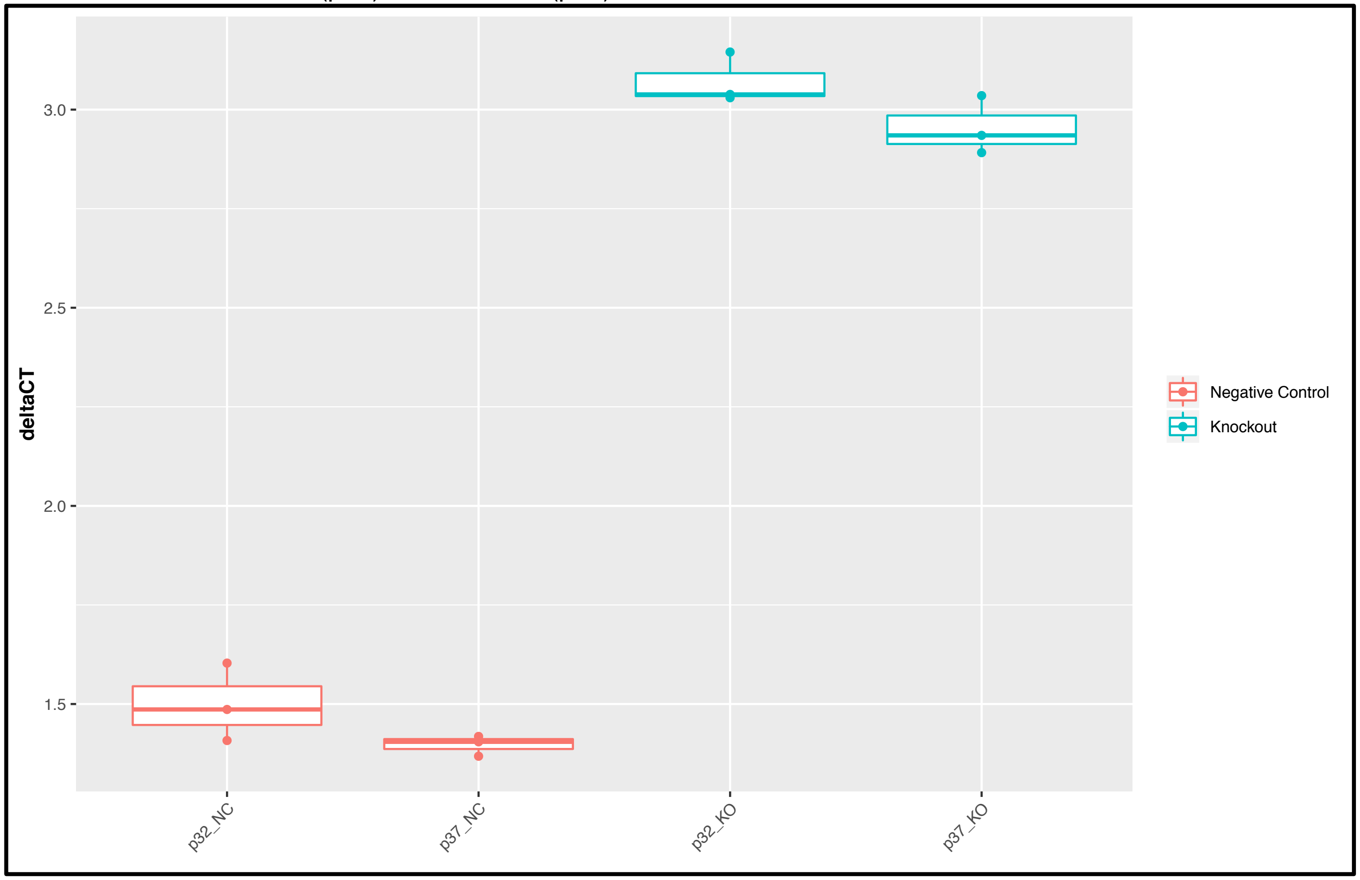

**Figure S8. ARIC identified CpGs with methylation differences between EPIC Negative Control (NC) lines and TFAM knockout (CRISPR) cell lines.** Units are methylation betas. *$P$<0.05.

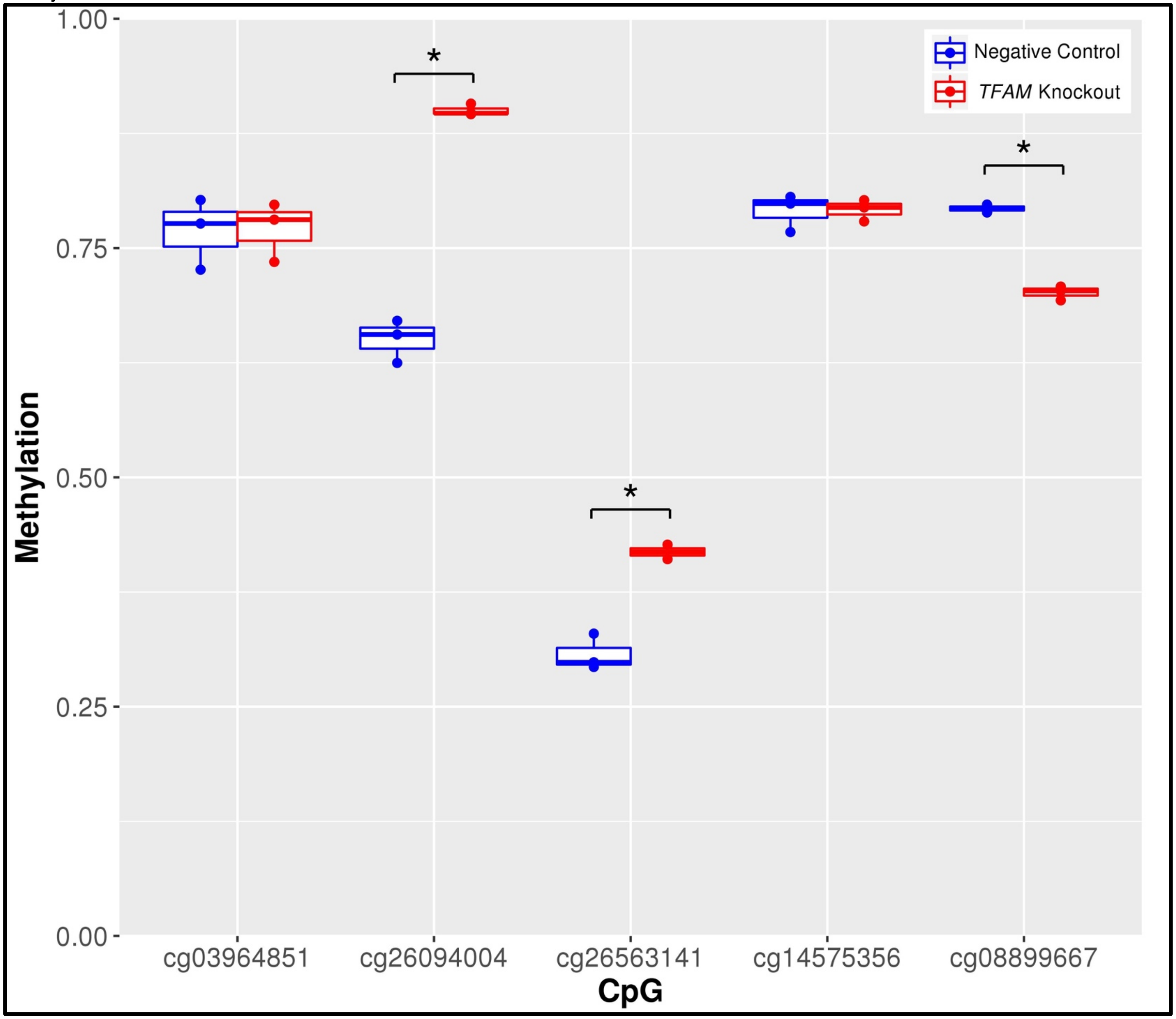

**Figure S9. Global methylation patterns in negative Control (NC) and CRISPR-TFAM knockout (CR) cell lines. A.** Average methylation of sample across all interrogated CpGs **B.** Density plot of methylation betas.

**A.**

| NC1 | NC2 | NC3 | CR1 | CR2 | CR3 |
|---|---|---|---|---|---|
| 0.513 | 0.505 | 0.522 | 0.510 | 0.499 | 0.510 |

**B.**

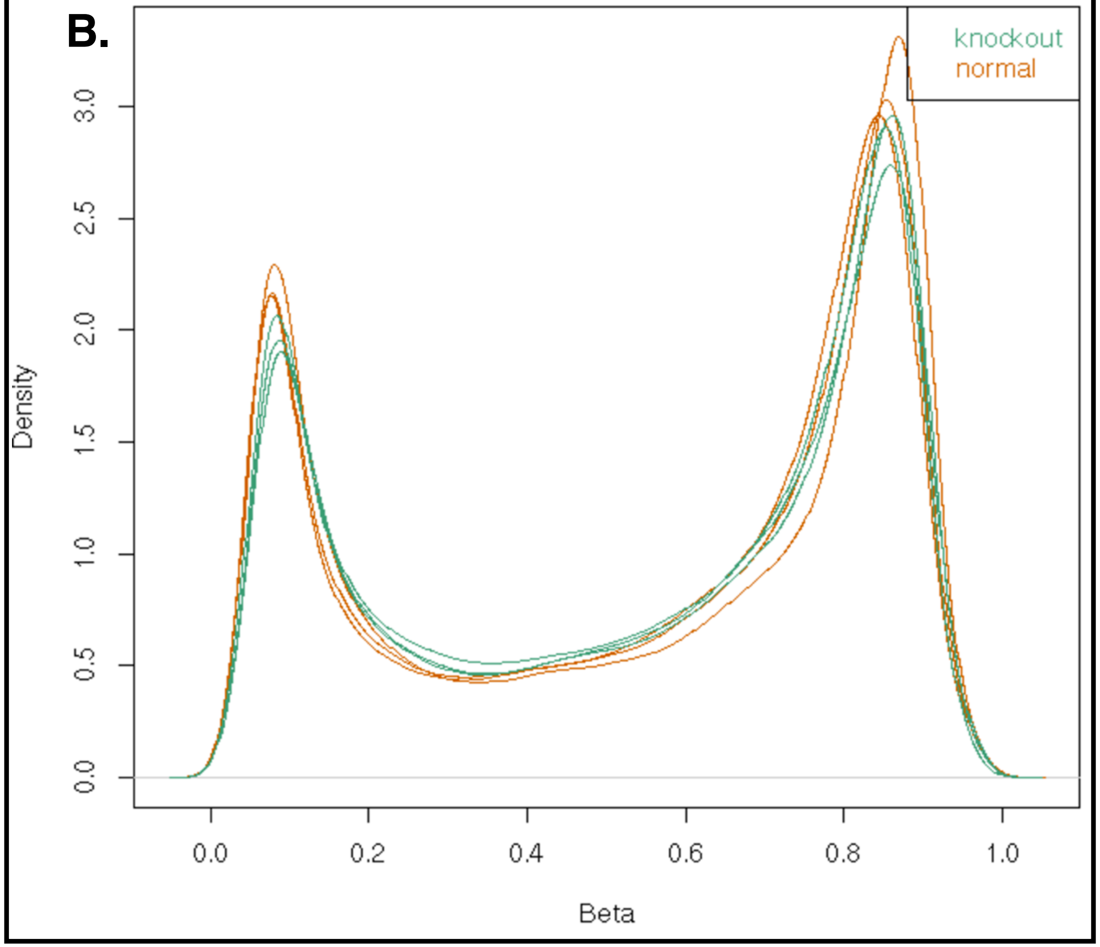

**Figure S10. TFAM knockout RNA sequencing clustering heatmap of TFAM knockout and control cell lines.** RNA sequencing was run in duplicate as indicated by a 1 or 2.

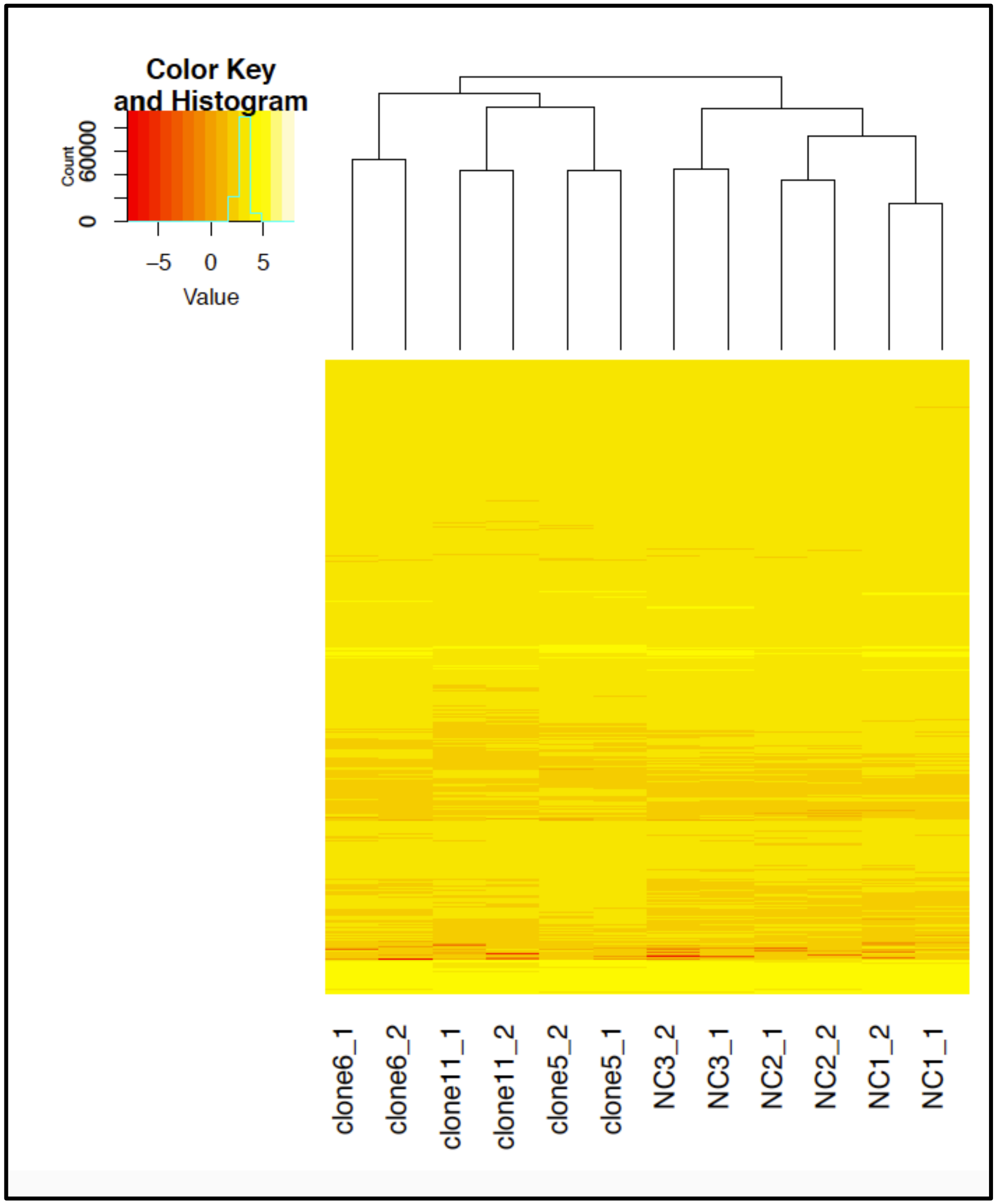
