## Supplemental Table S1 for "Mitochondrial DNA Copy Number (mtDNA-CN) Can Influence Mortality and Cardiovascular Disease via Methylation of Nuclear DNA CpGs"

**Table S1. Sample characteristics of discovery and validation cohorts.**

| | DISCOVERY COHORTS | | VALIDATION COHORTS | | | | |
|---|---|---|---|---|---|---|---|
| | ARIC AA (N=1567) | ARIC EA (N=940) | CHS | CHS AA (N=239) | CHS EA (N=294) | FHS | FHS (N=1995) |
| | *mean (range)* | *mean (range)* | | *mean (range)* | *mean (range)* | | *mean (range)* |
| **Age** | 57.2 (47 - 71) | 60.2 (47 - 72) | | 72.4 (65 - 92) | 72.3 (65 - 95) | | 61.4 (20-91) |
| **mtDNA-CN (SD units)** | 0.04 (-6.47 - 2.88) | 0.04 (-4.41 - 2.87) | | 0.05 (-3.19 - 2.69) | 0.05 (-2.28 - 2.76) | | 0.01 (-2.28 -8.11) |
| **Sex** | *N (percentage)* | *N (percentage)* | | *N (percentage)* | *N (percentage)* | | *N (percentage)* |
| Male | 604 (38.5%) | 381 (40.5%) | | 97 (41.8%) | 112 (38.9%) | | 901 (45.16%) |
| Female | 963 (61.5%) | 559 (59.5%) | | 135 (58.2%) | 176 (61.1%) | | 1094 (54.64%) |
| **Collection Site** | | | **Collection Site** | | | **Collection Site** | |
| Forsyth County, NC | 188 (12%) | 832 (88.5%) | Bowman Gray | 74 (31.9%) | 67 (23.3%) | Framingham Town | 1995 (100%) |
| Suburbs of Minneapolis, MN | 0 (0%) | 86 (9.2%) | Davis | 71 (30.6%) | 71 (24.7%) | | |
| Jackson, MS | 1379 (88%) | 0 (0%) | Hopkins | 0 (0%) | 83 (28.8%) | | |
| Washington County, MD | 0 (0%) | 22 (2.3%) | Pittsburgh | 87 (37.5%) | 67 (23.3%) | | |
| **Smoking Status** | | | | | | | |
| Current Smoker | 655 (41.8%) | 181 (19.3%) | | 32 (13.8%) | 26 (9.0%) | | 205 (10.4%) |
| Former Smoker | 484 (30.9%) | 366 (38.9%) | | 89 (38.4%) | 116 (40.3%) | | 947 (47.47%) |
| Never Smoker | 428 (27.3%) | 392 (41.7%) | | 85 (36.6%) | 133 (46.2%) | | 838 (42.0%) |
| Unknown | 0 (0.0%) | 1 (0.1%) | | 26 (11.2%) | 13 (4.5%) | | 5 (0.2%) |
| **Phenotypes (# of cases)** | | | | | | | |
| Mortality | 605 (38.6%) | 224 (23.8%) | | 194 (83.6%) | 263 (91.3%) | | 217 (10.87%) |
| ***CVD*** Prevalent | 154 (9.8%) | 49 (5.2%) | | N/A | N/A | | 94 (4.71%) |
| Incident | 296 (18.9%) | 108 (11.5%) | | 83 (35.8%) | 99 (34.4%) | | 94 (4.71%) |
| ***CHD*** Prevalent | 112 (7.1%) | 40 (4.3%) | | N/A | N/A | | 94 (4.71%) |
| Incident | 193 (12.3%) | 83 (8.8%) | | 48 (20.7%) | 57 (19.8%) | | 68 (3.41%) |
| **Cell Type Proportions** | *mean (range)* | *mean (range)* | | *mean (range)* | *mean (range)* | | *mean (range)* |
| CD8T Lymphocytes | 0.15 (0.00 - 0.48) | 0.10 (0.00 - 0.27) | | 0.09 (0.00 - 0.38) | 0.06 (0.00 - 0.22) | | 0.10 (0.00-0.36) |
| CD4T Lymphocytes | 0.19 (0.00 - 0.52) | 0.16 (0.00 - 0.44) | | 0.20 (0.00 - 0.48) | 0.15 (0.00 - 0.52) | | 0.19 (0.02-0.44) |
| B-cells | 0.07 (0.00 - 0.58) | 0.06 (0.00 - 0.56) | | 0.08 (0.00 - 0.26) | 0.06 (0.00 - 0.76) | | 0.04 (0.00-0.52) |
| Monocytes | 0.13 (0.02 - 0.26) | 0.09 (0.02 - 0.19) | | 0.10 (0.00 - 0.27) | 0.09 (0.01 - 0.35) | | 0.12 (0.05-0.30) |
| Granulocytes | 0.45 (0.15 - 0.98) | 0.55 (0.16 - 0.93) | | 0.44 (0.11 - 0.75) | 0.57 (0.03 - 0.92) | | 0.49 (0.02-0.85) |
| Natural Killer cells | N/A | 0.07 (0.00 - 0.36) | | 0.12 (0.01 - 0.38) | 0.09 (0.00 - 0.36) | | 0.02 (0.00-0.13) |

CVD: Cardiovascular disease. CHD: Coronary Heart Disease.
