## Supplemental Table S6 for "Mitochondrial DNA Copy Number (mtDNA-CN) Can Influence Mortality and Cardiovascular Disease via Methylation of Nuclear DNA CpGs"

**Table S6. Neuroactive-ligand receptor interaction genes as identified by KEGG analysis in each approach, number of genes used in each analysis and neuroactive ligand enrichment *P*-value are included in brackets.**

| ARIC Meta-Analysis (300 CpGs, $P$=5.24x10$^{-12}$) | *TFAM* Methylation (300 CpGs, $P$=4.41x10$^{-4}$) | *TFAM* Expression (169 genes, $P$=4.30x10$^{-4}$) | *TFAM* Integrated (Methylation/Expression) (188 genes, $P$=8.77x10$^{-6}$) |
|---|---|---|---|
| *CHRM2* | *GABBR1* | *GABRB1* | *GABRB1* |
| *CHRM3* | *GRIN3B* | *MC2R* | *MC2R* |
| *CTSG* | *GABRA5* | *GH1* | *GH1* |
| *AGTR1* | *GABRB1* | *GABRA2* | *GABRA2* |
| *GABRG3* | *GABRG3* | *GABRG1* | *GABRG1* |
| *GHR* | *GABRB3* | *ADRB2* | |
| *GRIA2* | *GALR1* | *MC4R* | |
| *GRIA4* | *NPBWR1* | | |
| *P2RX1* | *GRIK1* | | |
| *P2RY2* | *GRIN2D* | | |
| *PTGER2* | *HTR1E* | | |
| *HTR1B* | *TRHR* | | |
| *NTSR1* | *TSHR* | | |
| | *VIPR2* | | |
| | *CCKBR* | | |
