## Supplemental Table S7 for "Mitochondrial DNA Copy Number (mtDNA-CN) Can Influence Mortality and Cardiovascular Disease via Methylation of Nuclear DNA CpGs"

**Table S7. Methylation Status of Validated CpGs in *TFAM* KO cell lines (N=6).** Bolded entries indicate differential expression *P*<0.05.

| Marker Name | All Cohort Meta-Analysis | | | | Average Methylation in Negative Control Lines | Average Methylation in *TFAM* Knockout Lines | *TFAM* Differential Expression | |
|---|---|---|---|---|---|---|---|---|
| | Mean Methylation | Estimate | Standard Error | *P*-value | | | Beta Estimate | *P*-Value |
| cg03964851 (surrogate for cg21051031) | 0.83 | 0.0038 | 0.0004 | 7.34E-27 | 0.7685 | 0.7710 | -0.0025 | 9.42E-01 |
| **cg26094004** | 0.55 | -0.0079 | 0.0007 | 4.13E-28 | 0.6504 | 0.9001 | -0.2497 | 2.91E-05 |
| **cg26563141** | 0.37 | -0.0060 | 0.0008 | 2.20E-14 | 0.3071 | 0.4187 | -0.1116 | 1.25E-02 |
| cg14575356 | 0.55 | 0.0033 | 0.0005 | 1.22E-09 | 0.7906 | 0.7918 | -0.0013 | 9.40E-01 |
| cg23513930 | 0.35 | 0.0020 | 0.0003 | 3.71E-09 | Not on EPIC array and no surrogate available | | | |
| **cg08899667** | 0.58 | -0.0041 | 0.0006 | 1.55E-12 | 0.7931 | 0.7014 | 0.0917 | 3.33E-03 |

***Note:** Mean methylation for cg21051031 = 0.85
