## Supplemental Table S8 for "Mitochondrial DNA Copy Number (mtDNA-CN) Can Influence Mortality and Cardiovascular Disease via Methylation of Nuclear DNA CpGs"

**Table S8. Differentially expressed genes (*P*<0.05) within 1 Mb of differentially methylated CpGs in *TFAM* knockout cell lines. Shading indicates most differentially expressed gene for each CpG.**

| EWAS CpG | Chr: Position | Number of Genes Within 1 Mb | *P*-Value for *TFAM* Methylation Difference | Gene | Test Statistic for *TFAM* Expression | *P*-value for *TFAM* Expression | Direction of Effect (following KO)* | Distance from CpG (Kb) | Description |
|---|---|---|---|---|---|---|---|---|---|
| **cg26094004** | **17: 42,075,116** | **42** | 2.91E-05 | *ACLY* | 4.9871 | 2.55E-02 | Negative | 144.6 | ATP citrate lyase [Source:HGNC Symbol;Acc:HGNC:115] |
| | | | | *KAT2A* | 5.0487 | 2.46E-02 | Negative | 38.0 | lysine acetyltransferase 2A [Source:HGNC Symbol;Acc:HGNC:4201] |
| | | | | *HSPB9* | 8.3116 | 3.94E-03 | Negative | 46.3 | heat shock protein family B (small) member 9 [Source:HGNC Symbol;Acc:HGNC:30589] |
| | | | | *KCNH4* | 9.5817 | 1.97E-03 | Negative | 81.8 | potassium voltage-gated channel subfamily H member 4 [Source:HGNC Symbol;Acc:HGNC:6253] |
| | | | | *COASY* | 4.3154 | 3.78E-02 | Negative | 486.4 | Coenzyme A synthase [Source:HGNC Symbol;Acc:HGNC:29932] |
| | | | | *CCR10* | 4.5647 | 3.26E-02 | Positive | 603.8 | C-C motif chemokine receptor 10 [Source:HGNC Symbol;Acc:HGNC:4474] |
| | | | | *RAMP2* | 11.9339 | 5.51E-04 | Positive | 683.3 | receptor activity modifying protein 2 [Source:HGNC Symbol;Acc:HGNC:9844] |
| | | | | *AOC3* | 4.2112 | 4.02E-02 | Positive | 776.1 | copper containing 3 [Source:HGNC Symbol;Acc:HGNC:550] |
| | | | | *IFI35* | 16.9886 | 3.76E-05 | Positive | 931.6 | interferon induced protein 35 [Source:HGNC Symbol;Acc:HGNC:5399] |
| | | | | *RND2* | 8.9365 | 2.80E-03 | Positive | 950.1 | Rho family GTPase 2 [Source:HGNC Symbol;Acc:HGNC:18315] |
| | | | | *BRCA1* | 4.0986 | 4.29E-02 | Positive | 969.2 | DNA repair associated [Source:HGNC Symbol;Acc:HGNC:1100] |
| **cg26563141** | **2: 88,124,876** | **4** | 1.25E-02 | *RPIA* | 20.8215 | 5.04E-06 | Negative | 566.8 | ribose 5-phosphate isomerase A [Source:HGNC Symbol;Acc:HGNC:10297] |
| **cg08899667** | **6: 31,761,055** | **32** | 3.33E-03 | *CCHCR1* | 8.0190 | 4.63E-03 | Positive | 602.8 | coiled-coil alpha-helical rod protein 1 [Source:HGNC Symbol;Acc:HGNC:13930] |
| | | | | *HLA-C* | 4.6442 | 3.12E-02 | Positive | 488.9 | major histocompatibility complex: class I: C [Source:HGNC Symbol;Acc:HGNC:4933] |
| | | | | *HLA-B* | 9.4318 | 2.13E-03 | Positive | 403.9 | major histocompatibility complex: class I: B [Source:HGNC Symbol;Acc:HGNC:4932] |
| | | | | *ATP6V1G2* | 7.1309 | 7.58E-03 | Positive | 212.6 | ATPase H+ transporting V1 subunit G2 [Source:HGNC Symbol;Acc:HGNC:862] |
| | | | | *NFKBIL1* | 5.4633 | 1.94E-02 | Positive | 202.2 | NFKB inhibitor like 1 [Source:HGNC Symbol;Acc:HGNC:7800] |
| | | | | *CLIC1* | 4.2279 | 3.98E-02 | Positive | 21.3 | chloride intracellular channel 1 [Source:HGNC Symbol;Acc:HGNC:2062] |
| | | | | *MSH5* | 13.4158 | 2.50E-04 | Positive | 1.8 | mutS homolog 5 [Source:HGNC Symbol;Acc:HGNC:7328] |
| | | | | *HSPA1L* | 4.7247 | 2.97E-02 | Positive | 48.6 | heat shock protein family A (Hsp70) member 1 like [Source:HGNC Symbol;Acc:HGNC:5234] |
| | | | | *C2* | 5.9338 | 1.49E-02 | Positive | 136.7 | complement C2 [Source:HGNC Symbol;Acc:HGNC:1248] |
| | | | | *FKBPL* | 8.1388 | 4.33E-03 | Positive | 367.7 | FK506 binding protein like [Source:HGNC Symbol;Acc:HGNC:13949] |
| | | | | *EGFL8* | 4.5363 | 3.32E-02 | Negative | 403.5 | EGF like domain multiple 8 [Source:HGNC Symbol;Acc:HGNC:13944] |
| | | | | *HLA-DRA* | 10.6612 | 1.09E-03 | Positive | 678.8 | major histocompatibility complex: class II: DR alpha [Source:HGNC Symbol;Acc:HGNC:4947] |
| | | | | *HLA-DRB5* | 24.7584 | 6.50E-07 | Negative | 756.3 | major histocompatibility complex: class II: DR beta 5 [Source:HGNC Symbol;Acc:HGNC:4953] |
| | | | | *HLA-DRB1* | 4.1996 | 4.04E-02 | Positive | 817.7 | major histocompatibility complex: class II: DR beta 1 [Source:HGNC Symbol;Acc:HGNC:4948] |

*Positive beta indicates that after the decrease in mtDNA-CN (*TFAM* Knockout), expression has increased as compared to controls.
