## Supplemental Table S9 for "Mitochondrial DNA Copy Number (mtDNA-CN) Can Influence Mortality and Cardiovascular Disease via Methylation of Nuclear DNA CpGs"

**Table S9. Results of Mendelian Randomization. A.** Results for association between ARIC EA and AA derived independent *cis* meQTLs and mtDNA-CN. **B.** Results for association between ARIC meta-analysis derived independent *cis* meQTLs and mtDNA-CN (fixed effects model).

**A.**

| ARIC Cohort | Chr | meQTL CpG from EWAS | CpG Position | meQTL SNP | SNP Position | MAF | Imputation Quality ($R^2$) | mtDNA~meQTL SNP | | | CpG~meQTL SNP | | | Power for MR |
|---|---|---|---|---|---|---|---|---|---|---|---|---|---|---|
| | | | | | | | | Beta Estimate | Standard Error | *P*-value | Beta Estimate | Standard Error | *P*-value | |
| EA (Permuted *P*=7.84E-04) | 17 | cg26094004 | 42,075,116 | rs11654132 | 42,149,134 | 0.178 | 0.97 | -0.0327 | 0.0779 | 0.67 | 0.0322 | 0.0043 | 8.80E-14 | 0.99 |
| | 6 | cg08899667 | 31,761,055 | rs3117574 | 31,725,230 | 0.098 | 1.00 | 0.0279 | 0.0704 | 0.69 | 0.0149 | 0.0031 | 1.76E-06 | 0.29 |
| | 6 | cg08899667 | 31,761,055 | rs9267653 | 31,840,415 | 0.305 | 0.97 | -0.0126 | 0.0507 | 0.80 | 0.0101 | 0.0022 | 6.42E-06 | 0.28 |
| | 5 | cg21051031 | 93,905,482 | rs2973154 | 93,930,488 | 0.195 | 0.97 | 0.0115 | 0.0615 | 0.85 | 0.0073 | 0.0021 | 4.47E-04 | 0.75 |
| AA (Permuted *P*=9.12E-04) | 3 | cg23513930 | 10,334,717 | rs154236 | 10,356,314 | 0.254 | 0.96 | -0.0724 | 0.0417 | 0.08 | -0.0064 | 0.0009 | 2.23E-11 | 0.86 |
| | 6 | cg08899667 | 31,761,055 | rs28366163 | 31,704,804 | 0.148 | 0.94 | 0.0114 | 0.0474 | 0.81 | 0.0097 | 0.0019 | 4.64E-07 | 0.75 |
| | 6 | cg14575356 | 130,013,903 | rs1894642 | 130,015,331 | 0.410 | 1.00 | 0.0375 | 0.0354 | 0.29 | -0.0056 | 0.0012 | 3.95E-06 | 0.58 |
| | 6 | cg14575356 | 130,013,903 | rs17469966 | 130,061,044 | 0.057 | 0.82 | 0.1041 | 0.0840 | 0.22 | -0.0129 | 0.0030 | 1.70E-05 | 0.49 |
| | 6 | cg08899667 | 31,761,055 | rs9267659 | 31,846,234 | 0.057 | 0.98 | 0.0483 | 0.0856 | 0.57 | 0.0151 | 0.0036 | 2.52E-05 | 0.56 |
| | 6 | cg14575356 | 130,013,903 | rs9398917 | 130,014,226 | 0.238 | 0.98 | -0.0272 | 0.0391 | 0.49 | -0.0046 | 0.0014 | 6.44E-04 | 0.34 |

**B.**

| Chr | meQTL CpG from EWAS | CpG Position | meQTL SNP | SNP Position | Analysis | MAF | Imputation Quality ($R^2$) | mtDNA~meQTL SNP | | | CpG~meQTL SNP | | | Power for MR |
|---|---|---|---|---|---|---|---|---|---|---|---|---|---|---|
| | | | | | | | | Beta Estimate | Standard Error | *P*-value | Beta Estimate | Standard Error | *P*-value | |
| 6 | cg08899667 | 31,761,055 | rs9267653 | 31,840,415 | Meta | | | -0.0186 | 0.0309 | 0.55 | 0.0086 | 0.0013 | 1.99E-11 | |
| | | | | | EA | 0.305 | 0.97 | -0.0126 | 0.0507 | 0.80 | 0.0101 | 0.0022 | 6.42E-06 | 0.28 |
| | | | | | AA | 0.246 | 0.97 | -0.0221 | 0.0390 | 0.57 | 0.0078 | 0.0015 | 5.31E-07 | 0.75 |
| 6 | cg08899667 | 31,761,055 | rs28366163 | 31,704,804 | Meta | | | 0.0335 | 0.0425 | 0.43 | 0.0107 | 0.0017 | 1.10E-09 | |
| | | | | | EA | 0.063 | 0.96 | 0.1234 | 0.0957 | 0.20 | 0.0143 | 0.0041 | 4.54E-04 | 0.18 |
| | | | | | AA | 0.148 | 0.94 | 0.0114 | 0.0474 | 0.81 | 0.0096 | 0.0019 | 4.18E-07 | 0.75 |
